# Inhibiting LSD1 suppresses coronavirus-induced inflammation but spares innate antiviral activity

**DOI:** 10.1101/2021.05.02.441948

**Authors:** Luca Mazzarella, Fabio Santoro, Roberto Ravasio, Paul E. Massa, Simona Rodighiero, Elena Gavilán, Mauro Romanenghi, Bruno Achutti Duso, Emanuele Bonetti, Rani Pallavi, Deborah Trastulli, Isabella Pallavicini, Claudia Gentile, Tommaso Leonardi, Sebastiano Pasqualato, Gabriele Buttinelli, Angela Di Martino, Giorgio Fedele, Ilaria Schiavoni, Paola Stefanelli, Giuseppe Meroni, Christian Steinkuhler, Gianluca Fossati, Saverio Minucci, Pier Giuseppe Pelicci

## Abstract

Tissue-resident macrophages exert critical but conflicting effects on the progression of coronavirus infections by secreting both anti-viral type I Interferons and tissue-damaging inflammatory cytokines. Steroids, the only class of host-targeting drugs approved for Covid19, indiscriminately suppress both responses, possibly impairing viral clearance, and provide limited clinical benefit. Here we set up a mouse *in vitro* co-culture system that reproduces the macrophage response to SARS-CoV2 seen in patients and allows quantitation of inflammatory and antiviral activities. We show that the NFKB-dependent inflammatory response can be selectively inhibited by ablating the lysine-demethylase LSD1, which additionally unleashed interferon-independent ISG activation and blocked viral egress through the lysosomal pathway. These results provide a rationale for repurposing LSD1 inhibitors, a class of drugs extensively studied in oncology, for Covid-19 treatment.

**One-Sentence Summary:** Targeting a chromatin-modifying enzyme in coronavirus infections curbs tissue-damage without affecting antiviral response

## Introduction

Increasing evidence suggests that the severity of Covid-19 pathology is dictated by the profile of immune mediators produced in response to SARS-CoV2 infection. High levels of inflammatory cytokines typical of the innate immune response, such as IL6, TNF*α* and IL1, are associated with severe disease, whereas expression of molecules with direct antiviral activity, such as type I interferon and its downstream targets (Interferon-Stimulated Genes, ISGs) are associated with better outcome ^1–3^. Key transcriptional regulators of these innate responses to pathogens include Nuclear Factor-Kappa B (NF-κB), which regulates transcription of proinflammatory cytokines, and the family of Interferon-Regulated Factors (IRFs), which promote expression of Interferons and ISGs ^4^. These responses are structured in self-amplifying paracrine loops where downstream targets (e.g. IL1β and TNF*α* for NF-κB; type I Interferons for IRFs) are themselves activators of the same pathway, a logic architecture that allows the signal to propagate in space and time to rapidly arrest infection spread in the involved organ ^5, 6^. Unbridled loop activation, however, can lead to extensive local or systemic damage, further amplified by the recruitment of additional effector cells like neutrophils or lymphocytes.

The initiation and duration of the innate transcriptional responses is further regulated by multiple cofactors, among which chromatin-modifying enzymes are thought to play a key role ^7^. The Lysine Demethylase 1 (LSD1; also known as KDM1A), in particular, has been implicated in both NF-κB and IRF pathways and is pharmacologically targetable. Multiple LSD1 inhibitors have completed initial phases of clinical development for oncological indications with acceptable toxicity profiles ^8^ and may be amenable to repurposing. In a mouse system of Lipopolysaccharide (LPS)-mediated NF-κB activation using bone marrow-derived macrophages (BMDM), genetic ablation of *Lsd1* was found to be associated with decreased nuclear translocation of NF-ΚB and reduced binding to its target promoters, resulting in decreased transcription of proinflammatory cytokines and improved survival in a sepsis mouse model ^9^. On the other hand, LSD1 has also been implicated in suppressing endogenous retroviral elements (ERVs) in mouse melanoma cells. Loss of LSD1 induced up-regulation of ERV expression with consequent accumulation of double-stranded RNA (dsRNA), recognition by the dsRNA-sensing pathway MDA5-IRF3 and activation of an interferon-mediated response ^10^.

Transcription factor dynamics in human and murine coronavirus infections have been mostly assessed using epithelial or fibroblast cell lines ^11, 12^. Although epithelia and connective tissue are sites of active involvement in the pathogenesis of coronavirus infections, increasing evidence points to a central role for cells of the monocyte-macrophage compartment, which are strongly modulated during Covid-19 in relation to disease severity ^13–15^. These cells are equipped with a vast array of pathogen-sensing mechanisms and play a key role in the generation and amplification of inter-cellular signaling loops in innate immunity ^16–18^. Notably, alveolar macrophages from Covid-19 patients or SARS-CoV2-infected African Green Monkeys contain viral RNA in significant amounts, including negative strand sequences indicative of active replication ^19, 20^. However, the study of the molecular mechanisms involved in macrophage responses to human SARS-CoV2 has been hampered by the lack of *in vitro* models of productive infection ^21^, as also observed for SARS-CoV1 ^22, 23^, suggesting that virus uptake by macrophages is mediated by elements of the *in vivo* environment that are missing *in vitro*, such as non-receptor-mediated entry like antibody-mediated endocytosis or phagocytosis of infected cells ^20, 24^. In this study we investigated the dynamics of NF-κB and IRF activation in macrophage coronavirus infections and the potential role of LSD1.

## Results

### A system to separate macrophage-secreted cytotoxic and antiviral activities in response to coronavirus infection

As model system, we used mono- or co-cultures of mouse macrophages and fibroblasts or lung-epithelial cells infected with the Murine Hepatitis Virus (MHV), strain A59. MHV is a beta-coronavirus phylogenetically close to human SARS-CoV1/2 ^4^ and able to produce a disease highly similar to that triggered by human SARS-CoV ^25, 26^. MHV entry is mediated by the murine CEACAM1 receptor, which is highly expressed in mouse macrophages, both *in vivo* and *in vitro* ^27^, thus allowing to circumvent the intrinsic difficulties to model macrophage infection *in vitro* by SARS-CoVs. To investigate cell-intrinsic responses and paracrine signaling, bone-marrow derived macrophages (BMDM) were studied alone or in cocultures with fibroblasts (L929), an established model for the study of coronavirus innate responses ^12^ or the lung epithelial cell line LA4, previously shown to be susceptible to MHV-A59 infection ^28^ and chosen to confirm results on a model relevant for airborne infections. To measure extrinsic (secreted) and intrinsic responses to MHV infections, we set up a live-cell imaging system to quantitate cell death (by PI staining) and syncytia formation of H2BGFP-expressing L929 fibroblasts (L929^GFP^). This system (figure 1A), extensively described in the supplementary note, showed that UV-inactivated supernatants from MHV-infected BMDMs (snBMDM) contain distinct anti-viral and cytotoxic activities that can be quantified by measuring cell viability of L929^GFP^ and viral titers on L929 cells. We defined “extrinsic cytotoxic activity” (ECT) the loss of cell viability induced by snBMDM on uninfected L929^GFP^ cells (figure 1B) and “extrinsic antiviral activity” (EAV) the rescue of cell viability induced by snBMDM on infected L929^GFP^ cells (figure 1C). To verify if ECT and EAV can be biochemically separated, we applied progressive fractions of snBMDM obtained by size-exclusion chromatography to infected or uninfected L929^GFP^ cells and measured cell viability 48 hours post infection (hpi). As shown in figure 1D and F, the two activities were enriched in two clearly separated fractions, with ECT peaking in fractions 21-25 (corresponding to a predicted protein size of 50-60 kDa) and EAV peaking in fraction 27-31 (corresponding to a predicted protein size of ∼20 kDa). Previous studies identified TNF*α* and type I interferon as candidates for coronavirus-induced cytotoxic and antiviral activities respectively ^25, 29^. ELISA showed elevated levels of TNF*α* and IFN*α* in the fractions showing maximal ECT and EAV, respectively (figure 1E,G), consistent with their expected size (TNF*α* is biologically active in 52 kDa trimers ^24^). The role of TNF*α* and IFN*α* was confirmed in neutralization experiments with anti-TNF*α* anti-IFNAR or JAK inhibitors (figure 1H-K and supplementary figure 3A). Thus, MHV-activated macrophages secrete both anti-viral and cytototoxic activities, which can be easily dissected in our *in vitro* model and are mediated by the secretion of TNF*α* and IFN*α*, respectively.

**Figure 1.**
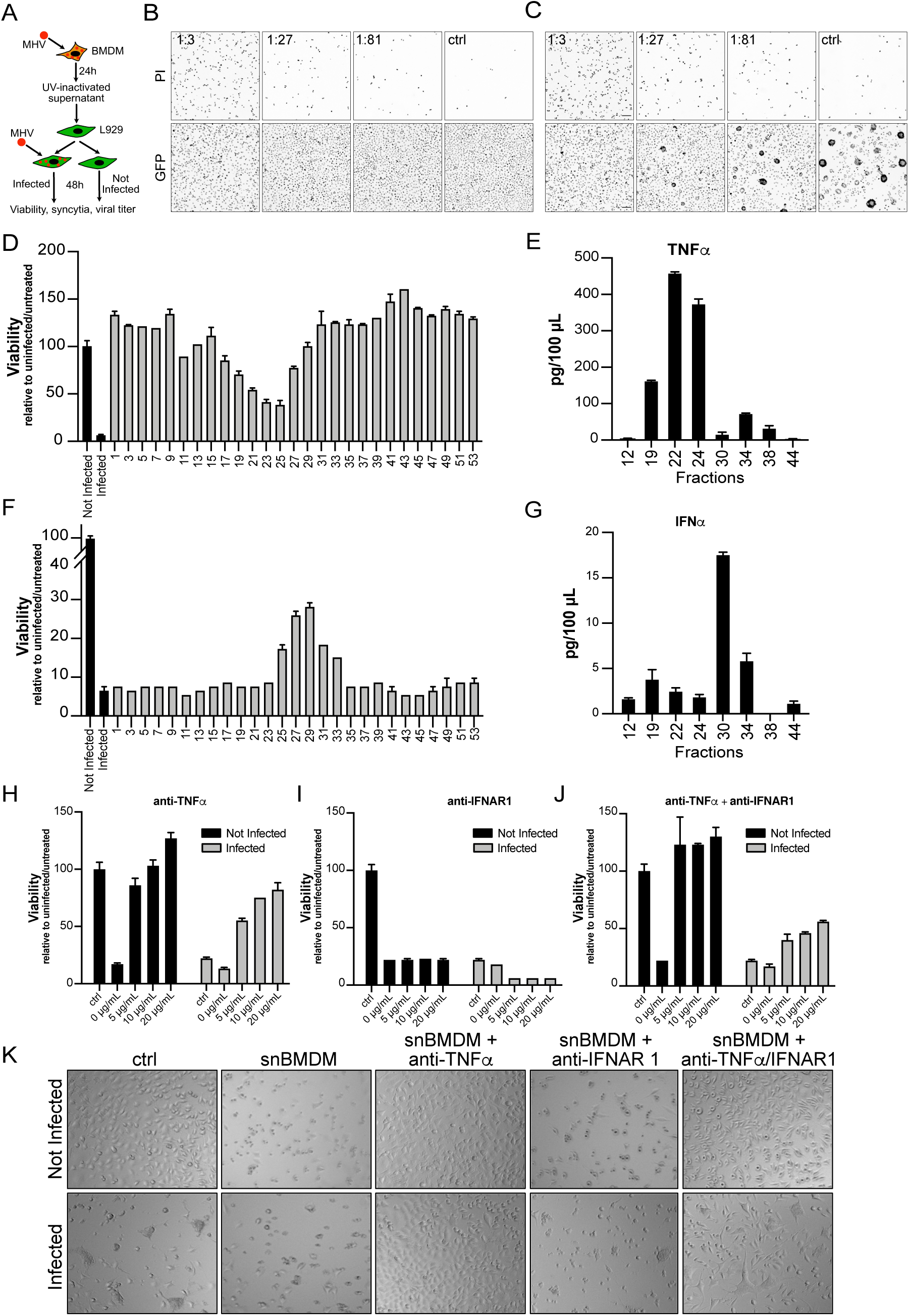
BMDM-secreted cytotoxic and antiviral activities are biochemically distinct and are sustained by TNF*α* and IFN*α*. **A**. Experimental strategy to evaluate BMDM extrinsic activity. **B,C**: Representative snapshots from time lapse microscopy imaging at 24 hours of L929^GFP^ cells not infected (B) or MHV infected at 0.1 MOI (C), exposed to MHV infected BMDM supernatants at the indicated dilutions. Scale bars: 100 μm. **D, F**: Effect of size-fractionated supernatant from MHV-infected (MOI 0.1) BMDMs on viability of non-infected (A) or infected MOI 0.1 (C) L929 cells. Viability is measured by Cell Titer Glo in triplicate (shown is mean + SD) and is expressed as fraction of uninfected and untreated cells. **E, G.** ELISA for TNF*α* and IFN*α* on size-fractionated supernatant from MHV-infected (MOI 0.1) BMDMs. Mean ±SD of triplicate wells per experiment. **H-J**. Effect of increasing doses of TNF*α* - or IFN*α* - neutralizing antibodies, alone or in combination, on 48h viability of non-infected or infected MOI 0.1 L929 cells exposed to supernatant from MOI 0.1-infected BMDM. Mean ±SD of triplicate wells per experiment. **K**. representative bright field images of non-infected or infected MOI 0.1 L929 cells 24 hours after treatment with control, supernatant from infected BMDM with or without TNF*α* - or IFN*α*-neutralizing antibodies, alone or in combination.

### Differential effect of LSD1 inhibition on macrophage-secreted cytotoxic vs antiviral activities

To test the role of LSD1 in coronavirus response, we tested the effect on EAV and ECT of treating BMDMs with either LSD1 inhibitor DDP38003 ^30^ (“snDDP”) or DMSO (“snDMSO”). DDP completely abrogated ECT, as viability of uninfected L929 cells was fully rescued (figure 2A, D) similar to what obtained with maximal doses of anti TNF*α* (figure 1H). Intruigingly, DDP did not reduce and actually enhanced EAV, as shown by better viability of infected L929 cells (figure 2B, E), and significantly reduced viral titers (figure 2C) and syncytia formation (figure 2F) compared to untreated cells. Size-exclusion chromatography confirmed that DDP 2.5μM completely eliminated ECT of fractions 21-25 (figure 2G) while maintaining EAV of fractions 27-31 (figure 2J); ELISA of snDDP confirmed abrogation of TNFa secretion (figure 2H,K) but persistence of IFNa (figure 2I,L). Neutralization experiments confirmed that enhanced EAV was sustained by persistent IFNa, as anti-IFNAR led to reactivation of the viral cytopathic effect (figure 2M). The role of LSD1 in ECT was genetically confirmed by LSD1 knockdown in the RAW264.7 macrophage cell line (supplementary figure 4F-G). Infected RAW264.7 secreted negligible EAV (no change in viability of infected L929 cells), but measurable ECT (80% viability loss in uninfected L929 cells, supplementary figure 4H). Consistent with DDP data, shLSD1 abrogated ECT and completely rescued viability of L929 cells exposed to infected RAW supernatant (supplementary figure 4H). Finally, to provide a minimal representation of the cellular landscape of *in vivo* coronavirus airborne infections, we also tested the impact of snBMDM on the lung epithelial cell line LA4, previously shown to be susceptible to MHV-A59 infection ^28^. LA4 were not sensitive to BMDM ECT but did indeed show loss of viability upon MHV infection, an effect that was partially reversed by snDDP (40% viability with snDDP 2.5 μM vs 20% in snDMSO, supplementary figure 4I).

**Figure 2.**
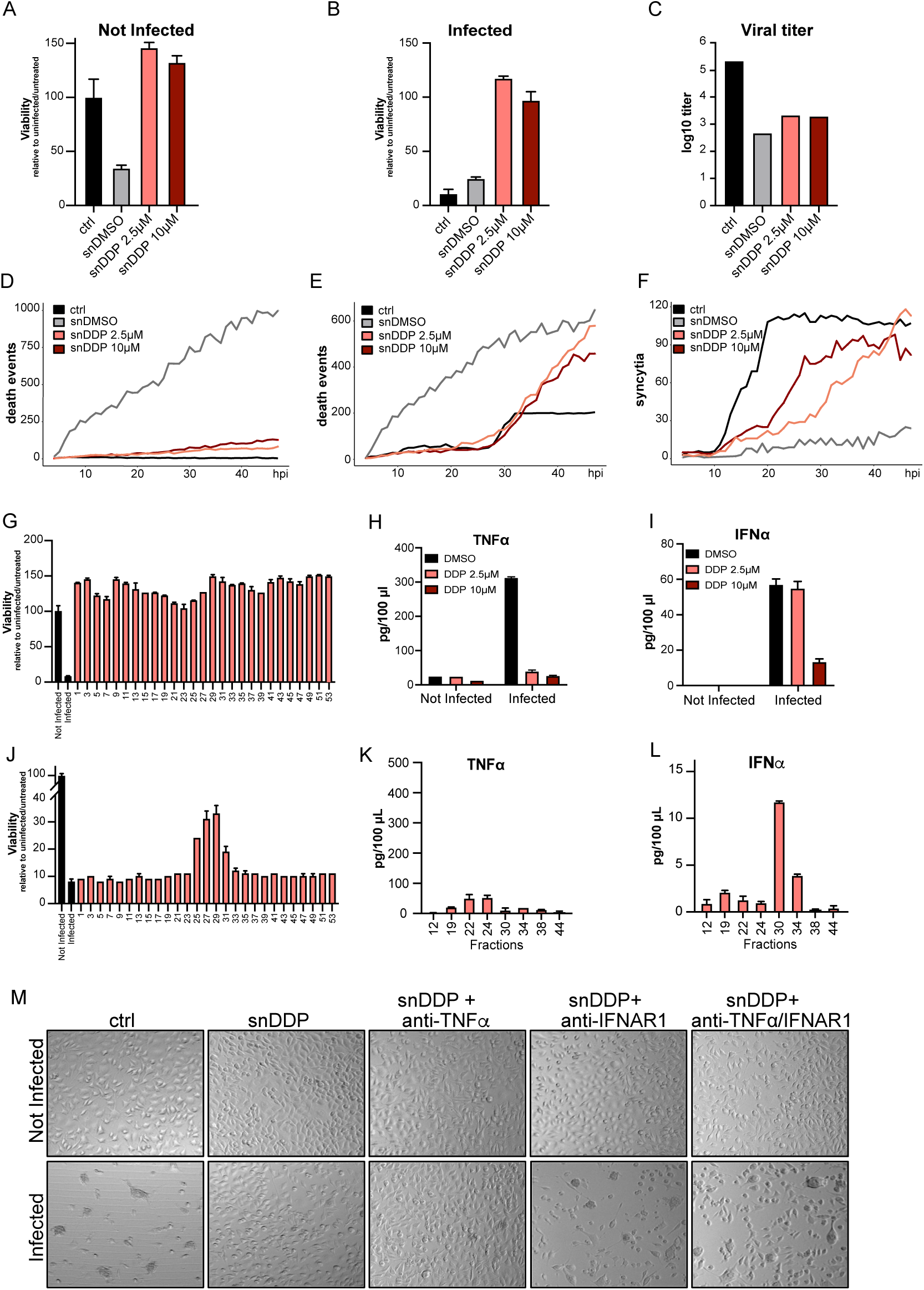
LSD1 inhibition ablates extrinsic cytotoxic activity but preserves extrinsic antiviral activity. **A-F.** Effect of LSD1 inhibitor DDP38003 on BMDM supernatant. BMDM were treated with vehicle (snDMSO) or LSD1 inhibitor DDP38003 (snDDP) at medium (2.5 μM) or high (10 μM) doses, 24 hours prior to infection with MHV-A59 at MOI 0.1. Supernatant was collected from BMDM cultures 24 hpi and applied to infected or uninfected L929 or L929^GFP^ cells. **A**: 48h viability of uninfected L929 cells measured by CTG; mean±SD of triplicate wells. **B**: 48h viability of infected L929 cells measured by CTG (for A and B mean and standard deviation of triplicate wells of one representative experiment are shown); mean±SD of triplicate wells. **C**. Viral titer measured by TCID50 24 hpi on L929 cells. **D** number of death events in time lapse imaging of uninfected L929^GFP^ cells. **E.** number of death events in time lapse imaging of infected L929^GFP^ cells. **F.** number of syncytia in time lapse imaging of infected L929^GFP^ cells live cell imaging. **G-J.** size-exclusion chromatography on supernatant from BMDM infected with MOI 0.1 and treated with DDP 2.5 μM. Biological activity of individual fractions was measured as in fig 2 on uninfected (G) or MOI 0.1-infected (J) L929 cells. **H-I**. ELISA for TNF*α* (H) and IFN*α* (I) on supernatant from BMDM infected or not with MHV 0.1 MOI and treated with vehicle (DMSO) or DDP at medium or high dosage. Mean ±SD of triplicate wells per experiment. **K,L.** ELISA for TNF*α* and IFN*α* on size-fractionated supernatant from BMDM infected with MHV 0.1 MOI and treated with vehicle (DMSO) or DDP at medium or high dosage. Mean ±SD of triplicate wells per experiment. **M** representative bright field images of non-infected or infected MOI 0.1 L929 cells 24 hours after treatment with control or supernatant from MOI 0.1-infected BMDM, treated with DDP 2.5 μM and treated with or without TNF*α* - or IFN*α* - neutralizing antibodies, alone or in combination.

To investigate molecular mechanisms underlying the macrophage response to MHV and the role of LSD1, we performed RNAseq analyses of macrophages treated with DMSO or DDP at moderate (2.5 μM) or high (10 μM) concentrations and infected or not with MHV-A59 MOI 0.1, at 24 hpi. Hierarchical clustering clearly separated infection and treatment groups, and revealed: i) a strong transcriptional effect of the viral infection; ii) a relatively modest impact of DDP on basal transcription; and iii) a strong and dose-dependent effect of DDP on MHV-dependent transcription (figure 3A and supplementary figure 5A). Seven transcriptional clusters were clearly demarcated, which were assigned to two groups, containing, respectively, genes up-(groups A1-4) or down- (groups B1-3) regulated by the infection. DDP largely antagonized the transcriptional effect of MHV-infection. In particular, in 5 of the 7 clusters, DDP markedly attenuated the down- (B2) or up- (A1,2) regulations induced by the infection, or even inverted their regulation (A3, B1). In only two clusters (A4, B3), instead, DDP intensified the up- (A4) or down- (B3) regulations induced by the infection (figure 3A, C). Gene ontology and motif-finding analyses revealed significant enrichment of distinct biological functions and transcription factors in each of the different clusters (figure 3B and supplementary table 2). As downregulation of the extrinsic cytotoxic activity is the dominant effect of DDP treatment, we initially focused on clusters containing genes whose MHV-dependent upregulation is inhibited by DDP (clusters A1-3). These clusters were enriched for proinflammatory cytokines and NF-ΚB binding motifs (cluster A1), Interferon-stimulated genes (ISG) and IRF binding motifs (cluster A2) and genes involved in granule formation and Sp1 binding motifs (cluster A3) (figure 3B). Cluster A1 included all cytokines currently implicated in the severe form of Covid-19 (*Il1a, I1b, Il6, Tnf*; supplementary table 3) and exhibited the strongest quantitative changes: it was the most highly upregulated in response to MHV (average of ∼8 fold) and the most downregulated in response to DDP (reaching baseline levels at 10 μM). The impact of DDP on IRF-associated genes in cluster A2 was significantly less pronounced: the extent of the MHV-induced upregulations in A2 was comparable to A1, but even with DDP 10 μM A2 cluster genes remained on average upregulated ∼5 fold (figure 3C). In further support of a direct activity of DDP on NF-ΚB, we observed extensive overlaps between genes of the A1-A4 clusters and the reported LPS-induced and LSD1-regulated NF-κB targets ^9^ (supplementary figure 5B).

**Figure 3.**
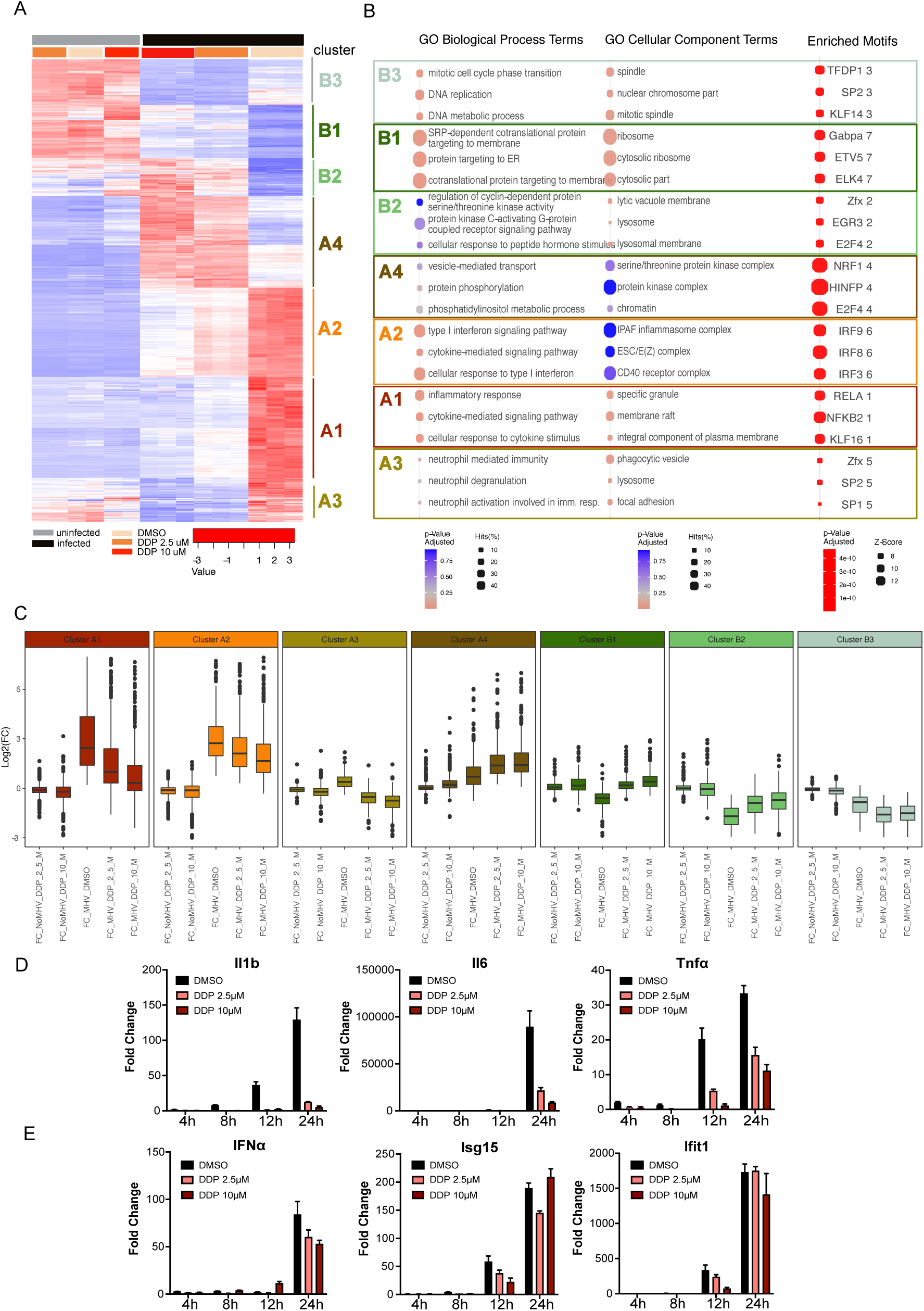
Effect of MHV infection and DDP treatment on macrophage transcriptional response. **A.** Hierarchical clustering of RNAseq data of BMDM infected or not with MHV-A59 MOI 0.1 and treated with DMS, DDP 2.5 μM or DDP 10 μM. Noninfected cells are in duplicate, infected in triplicate. **B.** Gene ontology and transcription factor binding motif enrichment by cluster. Top 3 gene sets per clusters are shown. Full list is in supplementary tables 2-3. **C.** boxplot of log2-fold changes of genes within each cluster over uninfected, untreated cells. Color code as in figures 4A,B. **D,E.** Gene expression analysis by qRT-PCR of representative NF-ΚB target genes (D) and IFN*α* and representative ISG (E) over the first 24 hpi in BMDM infected or not with MHV-A59 MOI 0.1 and treated with DMSO, DDP 2.5 μM or DDP 10 μM.

Gene expression analyses by qRT-PCR of representative NF-κB-dependent cytokines (Il1b, TNF*α*, Il6) showed rapid induction at 8-12 hpi and strong down-regulation by DDP (figure 3D). In turn, the impact of DDP on the expression of IFN*α* and representative ISGs (Isg15 and Ifit1) was significantly less pronounced (figure 3E). Notably, IFN*α* transcription was detected after that of Isg15 and Ifit1, suggesting an Interferon-independent initiation of ISG expression, a phenomenon previously reported in other viral infection systems ^31^.

LSD1 inhibition in cancer cells upregulates Transposable Elements (TE) and in particular Endogenous Retroviral Elements (ERVs), which lead to increased endogenous dsRNA and activation of the MDA5-IRF3 pathway ^10^. Thus, we assessed whether TE levels are affected by MHV infection and/or DDP treatment. We observed only a modest effect of DDP on TE RNAs in non-infected BMDMs (supplementary figure 6). TEs of multiple families including LINE, SINE and LTR were instead strongly upregulated upon MHV infection, in agreement with recent studies suggesting that coronavirus-dependent inflammation leads to TE upregulation ^32^. DDP dampened MHV-induced TE upregulation at both 2.5 and 10 μM concentrations (supplementary figure 6).

Activation of both NF-ΚB and IRFs is associated with nuclear localization ^5, 33^. Thus, we investigated levels of nuclear NF-ΚB and IRFs by biochemical fractionation and immunofluorescence. Western blotting analyses of nuclear/cytoplasmic fractions showed rapid nuclear re-localization of NF-ΚB after MHV-infection, as expected, already evident at 10 hpi (figure 6A-B). Same analysis for all 9 IRFs (supplementary figure 7A) showed upregulation and nuclear localization of IRF1 and, to a lesser extent, IRF2, whereas the other IRFs showed either no change (IRF5,9) or even decreased nuclear levels (IRF 3,4,6,7,8). Notable is the suppression of IRF3 and IRF7, typically involved in the response to dsRNA and ssRNA, thus confirming the existence of cellular-evasive mechanisms specific to coronaviruses ^12, 34, 35^. Results obtained by biochemical fractionation were confirmed by immunofluorescence analyses using anti-NF-ΚB and -IRF1 (figure 4C,D) antibodies, which showed nuclear localization of both factors upon MHV-infection. Notably, differential analyses of infected or uninfected cells (the former identified by NSP9 positivity) showed nuclear localization of both NF-ΚB and IRF1 also in NSP9-negative cells, suggesting activation of paracrine loops (figure 4C-D). Treatment of MHV-infected cells with DDP abrogated nuclear NF-ΚB but had a much less pronounced effect on IRF1, resulting in barely any difference at 24 hpi in fractionation experiments (Fig.4B) and a nuclear signal persistently above the baseline by immunofluorescence, both qualitatively and quantitatively (fig.4C,D). Chromatin Immunoprecipitation (ChIP) showed that MHV infection induced binding of both NF-ΚB and LSD1 at the promoter of NF-ΚB target genes, which was abrogated by DDP (figure 4E).

**Figure 4.**
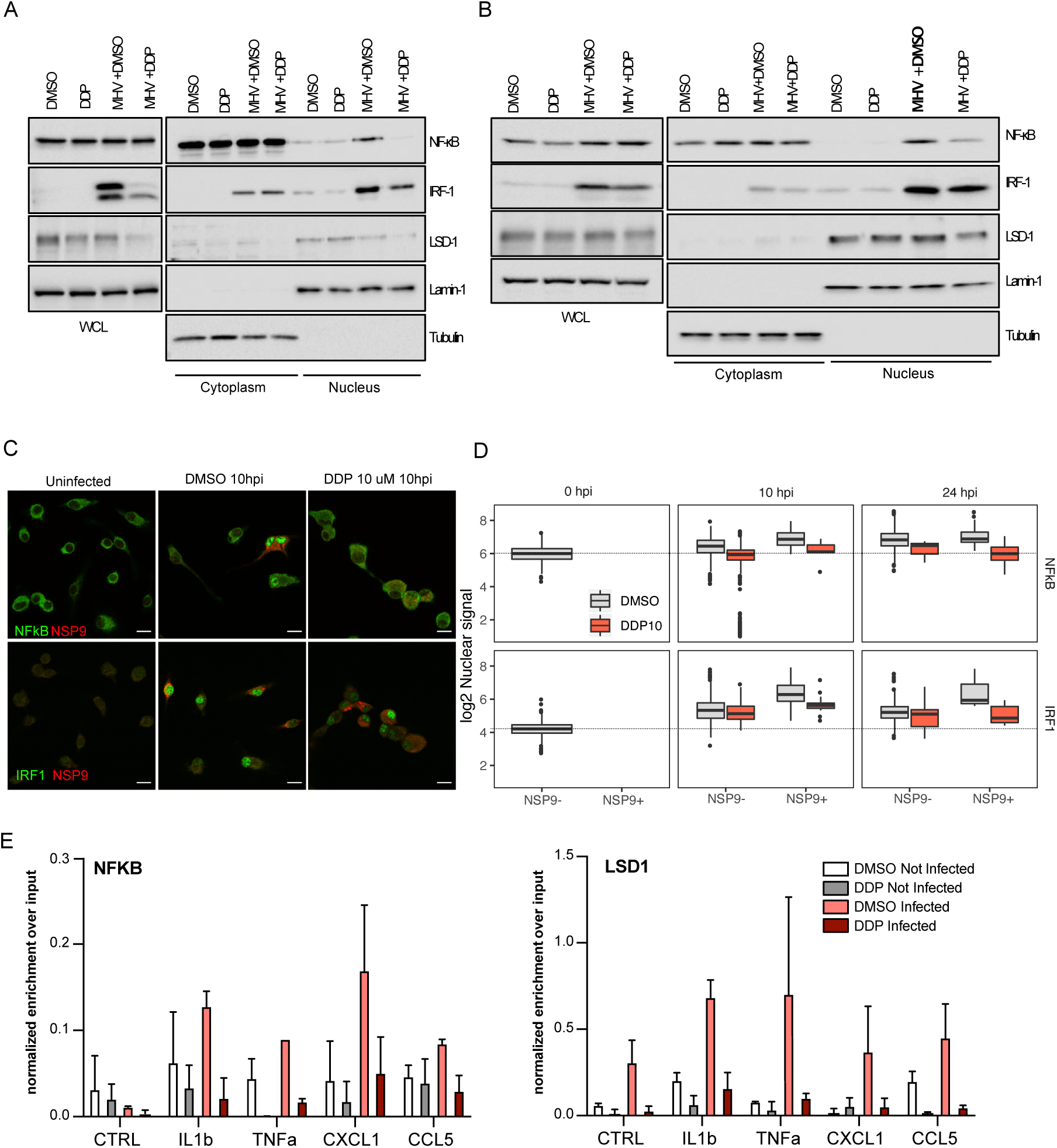
LSD1 inhibition abrogates NF-ΚB nuclear translocation but has only minimal effect on IRF1 nuclear translocation in response to MHV-A59 infection. **A-B** WB of whole cell lysate (WCL), cytoplasmic (C) and nuclear (N) fractions of BMDM at 10 hpi MOI 5 (**A**) and 24 hpi MOI 0.1 (**B**). **C, D**. IF for NF-ΚB and IRF1 showing subcellular localization in response to MHV; NSP9 identifies infected cells. In **C** representative images at 10 hpi MOI 5 are shown (scale bars: 10 μm). In **D**, boxplot of the mean nuclear signal for NF-ΚB (upper panel) or IRF1 (lower panel), divided by NSP9 positivity and treatment. Dashed line indicates the signal of uninfected and untreated cells at baseline. **E.** Chromatin IP for NF-ΚB and LSD1 in BMDM infected with MHV-A59 MOI 5 at 10 hpi, treated with DMSO or DDP 2.5 μM. Mean ± standard deviation of triplicate reactions.

Collectively, these results suggest that LSD1 is specifically implicated in regulating MHV-induced NF-ΚB nuclear relocalization and NFKB-dependent inflammatory cytokine production, leaving relatively unaltered the IRF1 response and the ensuing Interferon secretion.

### Effect of LSD1 inhibition on Interferon-independent ISG activation and lysosomal viral egress

In addition to the secreted (extrinsic) antiviral activity induced by DDP in MHV-infected BMDMs, we explored whether LSD1 inhibition also exerts a direct (intrinsic) antiviral activity, by measuring survival, titer and syncytia formation in L929, BMDMs and LA4 cells.

In all three cell types, treatment with DDP increased cell survival in a dose-dependent manner (figure 5A-B and supplementary figure 8) and reduced viral titer and syncytia formation (figure 6C-D and supplementary figure 7A, B, D, F). The magnitude of the antiviral activity was cell type-specific and most evident in L929 cells, where the titer was reduced by ∼1 log at 2.5 μM (figure 6C-D). In LA4 and BMDM, instead, a relevant effect was only evident at the higher dose of 10 μM (supplementary figure 8D,F). Importantly, LSD1 ablation by either CRISPR-Cas9 (figure 6E-F and supplementary figure 9A) or RNA interference (supplementary figure 9B) abrogated MHV-induced cytotoxicity and significantly reduced viral titer by at least 2 logs in L929 cells.

**Figure 5.**
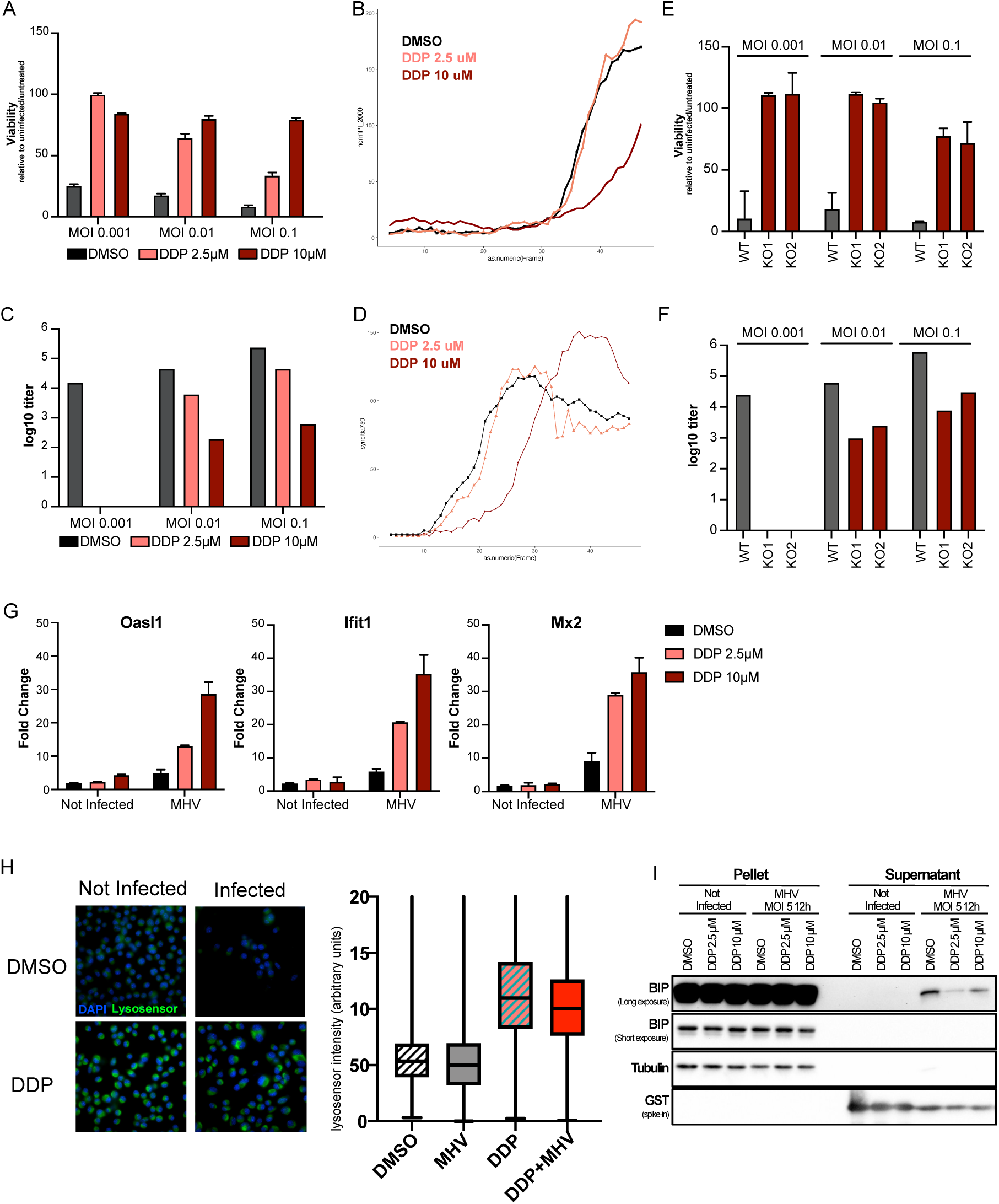
LSD1 inhibition exerts intrinsic antiviral activity and enhances lysosomal acidification. **A-D**. Direct antiviral effect on L929 cells, measured as survival by CTG (**A**), death events (**B**) and live cells (**C**) by live cell imaging, viral titer at 24 hpi (**D**) and syncytia formation by live cell imaging on L929^GFP^ cells. **E-F.** Effect of LSD1 KO (2 independent clones KO1 and KO2 vs non-targeted clone WT) on 48 hpi viability (E) or 24 hpi titer (F) of L929 cells infected with MHV at indicated MOI. **G.** Expression of selected ISGs by qRT-PCR in response to MHV-A59 MOI 0.1 and DMSO or DDP at increasing doses. **H.** Lysosome pH measurement using lysosensor on L929 cells uninfected or infected with MHV-A59 MOI 5 and treated with DMSO or DDP 10 μM, at 12 hpi. Representative images (left) and quantification (right). All distributions are significantly different by pairwise comparison with p values < 0.0001 by Mann-Whitney test (distributions are non-normal) and adjusted by Benjamini-Hochberg. **I.** BiP extrusion in L929 cells uninfected or infected with MHV-A59 MOI 5 and treated with DMSO or DDP 10 μM at 12 hpi, measured by WB on cell pellets and 1000 μl supernatant). The GST peptide was added as spike to the supernatant to equalize supernatant loading

We next investigated mechanisms of the direct anti-viral effect of LSD1 inhibition. L929 cells do not produce any type I interferon in response to MHV, irrespective of DDP treatment (supplementary figure 9C). Consistently, JAK inhibition did not antagonize the DDP protective effect (supplementary figure 9D). Of note, a subset of ISGs (*Ifit1, Oasl1, Mx2* among those tested) was modestly activated upon MHV and further upregulated by DDP in a dose-dependent manner (figure 5G), suggesting that ISG activation in DDP-treated L929 cells occurs in an Interferon-independent manner.

To further characterize the intrinsic antiviral activity of DDP we took advantage of BMDM RNAseq data, and focused on pathways enriched in clusters of the B group, which include genes that are either down-regulated or modestly affected by MHV, yet strongly upregulated by DDP treatment. These clusters were enriched for gene sets involved in vesicle biogenesis and lysosome biology (figure 3A-B). Coronavirus egress has been recently shown to occur through an unconventional lysosome-dependent pathway, which requires inhibition of lysosomal acidification and is associated with release of the ER marker BiP ^36^. In agreement, we observed downregulation by MHV and upregulation by DDP of the lysosomal *Atp6v1a* (supplementary figure 9E), a component of the vacuolar ATPase that mediates acidification of the organelle. In both BMDMs and L929 cells, DDP led to significant acidification of the lysosomal compartment, as measured by the pH-sensitive dye lysosensor, completely overriding the modest yet significant decrease in lysosomal acidification associated with MHV infection in L929 cells (figure 5H and supplementary figure 9F). Extracellular release of the endoplasmic reticulum chaperone BiP was reduced upon DDP treatment in both macrophages and L929 cells (figure 5I and supplementary figure 9G).

Together, these data suggest that LSD1 ablation allows the emergence of a cell-intrinsic antiviral response characterized by interferon-independent ISG activation and restoration of lysosomal acidification, resulting in reduced viral release. The precise point of regulation of this activity remains to be elucidated.

### LSD1 as a target to curb human cytokine storm in COVID-19 while preserving antiviral activity

To explore the translational relevance of LSD1 inhibition as a potential treatment for Covid-19 or other coronavirus infections, we compared the activity of DDP with ORY-1001 and dexamethasone. ORY-1001 is an LSD1 inhibitor of the same class of DDP that has completed initial phases of clinical development in the context of hematological malignances ^8^.

Dexamethasone is the only host-targeting medical treatment approved to date for Covid-19 as it moderately improves survival and effectively dampens the NF-κB-dependent response, but is also suspected to suppress interferon activity thus resulting in prolonged viral shedding ^37–39^. All three compounds inhibited the macrophage-secreted cytotoxic activity in a dose-dependent manner, as measured by restoration of cell viability of non-infected L929 cells exposed to BMDM-conditioned supernatants (figure 6A). When the same supernatants were added to infected L929 cells, even at the highest concentrations, neither of the two LSD1 inhibitors decreased the macrophage-secreted antiviral activity, as judged by restoration of cell viability and lack of syncytia formation. Dexamethasone, instead, at cytotoxic-suppressing dosages failed to recover viability and to prevent syncytia (10 μM, figure 6B,D). Similar results were obtained when the three drugs were applied directly on infected L929 cells: both DDP and Ory1001, but not dexamethasone, elicited intrinsic antiviral activity (figure 6C-D).

**Figure 6.**
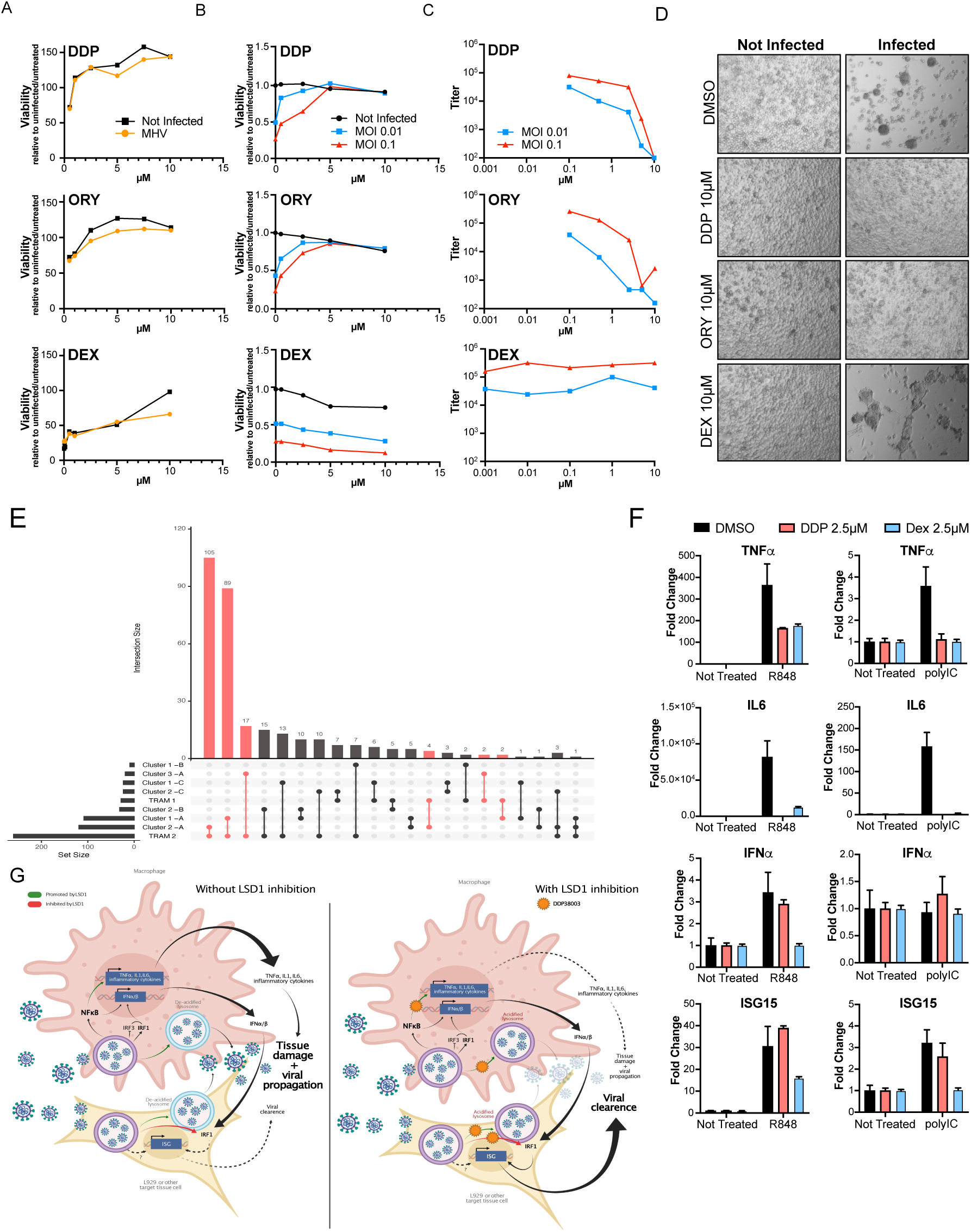
LSD1 inhibition is a viable target for human Covid-19 treatment. **A-D** response to equimolar doses of DDP38003 (DDP), Ory1001(ORY) and dexamethasone (DEXA) on macrophage extrinsic activity (A), cell survival of L929 cells and viral titer of L929 cells (C); representative bright field images of L929 cells infected at MOI 0.1 and treated as indicated are shown in D. **E.** Upset plot depicting the overlap of mouse BMDM transcriptional clusters (as in fig 3A) and human TRAM1/2 clusters from Querrey et al 2021. **F.** Expression level by qRT-PCR of NF-ΚB and ISGs in human macrophages stimulated with TLR3/RIGI/MDA5 agonist polyIC and TLR7/8 agonist R848. **G.** Model depicting innate responses to coronaviruses regulated by LSD1.

Then, we tested pharmacological interactions with type I Interferon, relevant for mechanistic understanding and potential pharmacological combinations. IFN*α* alone dose-dependently inhibited MHV activity and viral titer, and no synergism nor antagonism could be identified if cells were co-treated with increasing doses of DDP (supplementary figure 10A,B). Thus, although high-dose DDP does indeed reduce Interferon production by macrophages (figure 2I), it does not interfere with downstream interferon-dependent signaling.

As mentioned above, the possibility to directly study the response of human macrophages *in vitro* to SARS-CoV2 remains challenging. In our hands, as reported by others ^21^, attempts to infect monocyte-derived macrophages *in vitro* have been unsuccessful (supplementary figure 10C). Thus, we took advantage of the transcriptomic data from broncho-alveolar lavage of Covid-19 patients, recently published by Querrey et al ^20^, that show that in severe Covid-19 patients, Tissue-Resident Alveolar Macrophages infected by actively replicating SARS-CoV2 (TRAM2) have a transcriptional profile significantly different from that of from uninfected TRAMs (TRAM1) within the same patients. We found a striking overlap between TRAM2-overexpressed DEGs and genes of our clusters A-1 (NF-κB-enriched) and A-2 (IRF-enriched) (figure 6E and supplementary table 4). We compared the effect of DDP and dexamethasone on human monocyte-derived macrophages stimulated with agents inducing antiviral innate responses to dsRNA (polyIC) and ssRNA (R848). Again, whereas Dexamethasone led to generalized reduction of proinflammatory cytokines TNF*α* and IL6, DDP maintained selectivity with a significantly stronger reduction in the transcription of NF-ΚB-dependent cytokines than ISGs (figure 6F), demonstrating a general, trans-species and stimulus-independent selective effect of DDP on the inflammatory and interferon cascade. A general model for the role of LSD1 in regulating innate responses to coronaviruses is depicted in figure 6G.

## Discussion

Based on our findings we propose LSD1 inhibition as a promising strategy to prevent or treat Covid-19, with potential advantages over the only host-targeting drug class approved for Covid-19 to date, steroids, given the selective inhibition on NF-kB-dependent secretion of tissue-damaging proinflammatory with relative sparing or even enhancement of the Interferon response. The viability of this strategy is further supported by findings in murine models of SARS-CoV respiratory diseases, in which direct pharmacological inhibition of NF-ΚB or generation of viral mutants inducing attenuated NF-ΚB activation led to decreased disease severity and prolonged survival ^40^. In agreement with our findings, the GSK inhibitor GSK-LSD1 was recently shown to reduce levels of proinflammatory cytokines in peripheral blood leukocytes isolated from severe COVID-19 patients ^41^, although effect on tissue-resident macrophages and sparing of Interferon response were not shown. Oryzon has initiated the ESCAPE study with Vafidemstat in nonsevere Covid-19 (Eudract 2020-001618-39) but results are not yet known.

The need for cooperation between NF-κB and IRF factors in the so-called “enhanceosome” that regulates type I interferon expression ^42, 43^ may provide a mechanistic explanation for the differential activity of steroids *vs* LSD1 inhibitors: whereas steroids inhibit NF-ΚB by preventing NF-ΚB nuclear translocation through upregulation of IkB ^44^, LSD1 has been proposed to suppress the degradation of nuclear-translocated NF-ΚB ^9^. Thus, LSD1 inhibition may allow minute amounts of NF-ΚB to enter the nucleus for a restricted time window, allowing it to act as a “pioneer” transcription factor to render chromatin of the IRF1 target loci accessible for subsequent binding of canonical transcriptional activators ^45^. Further research is required to elucidate if histone demethylase activity is required for the effect of LSD1in the NF-κB response, and if main targets are NF-ΚB itself or critical effector proteins encoded by NF-κB target-genes.

An intriguing finding is the existence of an LSD1-dependent, interferon-independent cell-intrinsic response that results in the activation of a subset of ISGs and inhibition of lysosomal-mediated virus egress. The precise nature of this response, which may contribute to the initial, Interferon-independent trigger of IRF1, requires further investigation and may involve direct histone demethylation at regulatory elements of ISGs and/or genes involved in lysosomal activity. The finding that IRF1 and to a lesser extent IRF2 are the only IRFs that clearly change their intracellular localization in response to MHV infection is perhaps surprising, given the widespread involvement of other IRF members in the antiviral response to many viruses. However, although generally considered dispensable for the activation of antiviral responses^33^, IRF1 can lead to early activation of ISGs and type I Interferon, independently from other IRFs that typically occur as a later event, such as dsRNA-MDA5/RIG-I activated IRF3 ^31, 46, 47^. In the absence of other viral-induced effects, this would lead to Interferon-loop amplification, inhibition of TNF*α* signaling or IL1b production and attenuation of NF-κB signaling ^48, 49^. However, when IRF3-dependent amplification of type I Interferon is actively inhibited by coronavirus-specific mechanisms, cross-inhibition of the NF-κB loop is ineffective, thus favoring inflammation over Interferon-induced viral clearance as observed in severe Covid19 cases. Thus, sustaining interferon response while suppressing NF-kB through LSD1 inhibition may interrupt the propagation of the proinflammatory loop and allow viral clearance with minimal tissue damage, providing clinical benefit even in earlier disease stages in which steroids seem to be ineffective or even detrimental.

## Methods

### Data and materials availability

All unique/stable reagents generated in this study are available from the corresponding authors with a completed Materials Transfer Agreement.

The RNAseq dataset (fastq files, raw and normalized counts and differentially expressed genes) generated during this study are available at GEO GSE169399

The full sequence of the MHV-A59 strain used in the present study is deposited at NCBI (submission initiated, will be finalized upon acceptance)

### Viral stock

MHV strain A59 was kindly provided by Dr. Riccardo Villa at the IZSLER and propagated on L929 cells. Briefly, L929 cells were plated the day before infection at a density of 20 million cells in T175 flask. Cells were infected at MOI: 0,5 in 10 ml of free serum DMEM. After 1 hour incubation at 37 °C 40ml of DMEM supplemented with 3% FBS were added. Supernatant containing virus was harvested when the virus-induced cytopathic effect was visible on more than 70% of cells, usually 36 hours after infection. The identity of the virus was confirmed by Illumina sequencing (see supplementary table 1).

For SARS-CoV2 experiments, a strain isolated from an Italian patient in February 2020 ^50^ was used in the Biosafety level 3 facility of the Istituto Superiore di Sanita’ in Rome.

The virus was propagated in Vero E6 cells cultured at 37 °C in 5% CO_2_ in minimal essential medium (MEM) supplemented with 10% fetal calf serum (FCS), 1% l-glutamine, and 1.4% sodium bicarbonate. Virus-infected cells were maintained at 37 °C in 5% CO_2_ in MEM supplemented with 2% FCS. Titers were measured by CCID_50_ system in Vero E6 cell. Briefly, samples were serially diluted 1/10 in medium. Then 100 µL of each dilution was plated into ten wells of 96-well plates containing 80-90% confluent cells. The plates were incubated at 37 °C under 5% carbon dioxide for five days. Each well was then scored for the presence or absence of the virus. The limiting dilution end point (CCID_50_/ml) was determined by the Kärber equation.

### Cell culture

Raw264.7, L929, LA4, Calu3 and VeroE6 cell lines were purchased by American Type Culture Collection and grown according to ATCC recommendations. Cultures were maintained in a humidified tissue culture incubator at 37°C in 5% CO2. To assure mycoplasma-free conditions, all cells were routinely tested.

BMDM were obtained from bone marrow of 6-10 week old female C57Bl6 mice (Charles River). 10^6^ cells were plated in 10 cm untreated cell culture dishes, resuspended in 8 ml of Alpha MEM containing 20% FBS, 2 mM of L-glutamine, antibiotics, 40ng/ml of rm M-CSF (R&D System) and allowed to differentiate for 7 days.

Human monocytes were purified from peripheral blood collected from healthy blood donors at the Centro Trasfusionale Policlinico Umberto I, University La Sapienza blood bank (Rome, Italy) using Ficoll gradients (lympholyte-H; Cedarlane). CD14 cells were purified by anti-CD14 monoclonal antibody (mAb)-conjugated magnetic microbeads (Miltenyi Biotec) and then cultured for 6 days in RPMI 1640 medium (Life Technologies Invitrogen), supplemented with heat-inactivated 10% lipopolysaccharide-free FBS, 1 mM sodium pyruvate, 0.1 mM nonessential amino acids, 2 mM L-glutamine, 25 mM HEPES, 100 U/ml penicillin, 100 µg/ml streptomycin (all from EuroClone) in the presence of human recombinant M-CSF (100 ng/µL; Peprotech). Blood donors provided written informed consent for the collection of samples and subsequent analysis. Blood samples were processed anonymously.

### Generation of L929^GFP^ cells

L929 cells were plated at a density of 5X10^4^ in 24 well plate a day before the infection. A spin infection was used to infect L929 cells using a lentiviral vector carrying the H2B-GFP transgene ^51^. Briefly, concentrated H2B-GFP lentivirus (MOI 4) was added to L929 cell in a 24-well non-tissue culture-treated plate and centrifuge at 750 × *g*, 25 °C for 1 h. Infected cells were incubated for 3 h at 37 °C and replaced with fresh culture media (1 mL/well). After a recovery period of 2 days, GFP+ cells were sorted by fluorescence-activated cell sorting (FACS) and maintained in the culture for further experiments.

### *In vitro* pharmacological treatments

DDP38003 and Oryzon 1001 were synthesized as described in ^30^.

Dexamethasone (Sigma) was dissolved in DMSO.

PolyIC was purchased from Cytiva and dissolved in PBS to a concentration of 1mg/ml.

LPS was purchased from Sigma and dissolved in water to a concentration of 1mg/ml.

### Preparation of BMDM supernatant

BMDM were plated at a density of 500.000 cells per well in untreated cell culture 6 well plates, in a total volume of 2ml. After one night incubation, BMDM were treated with the indicated compounds. 24 hours later the cells were infected with MHV at the desired MOI, by replacing the overnight medium with DMEM 3% FCS containing MHV. After 1 hour, the viral inoculum was removed and replaced with BMDM medium supplemented with the drugs. 24 hours later, supernatant was harvested, clarified by centrifugation and used for subsequent experiments.

### UV inactivation of BMDM supernatant

500ul of BMDM supernatant were aliquoted in one 24 well and incubated on ice for 1 minute. UV inactivation was performed on ice, using Agilent Genomics/Stratagene Stratalinker 2400 UV Crosslinker, by delivering an energy dose equivalent to 0,3 Joules.

### Antiviral and cytotoxicity assay

L929 cells were plated at a density of 5000/well in 96 well plate, in a total volume of 100 µl. The day after, the culture medium was removed and replaced with 50 µl of the serially diluted UV inactivated BMDM supernatant. For TNFα and IFNAR neutralization assay, 50 µl of the serially diluted UV inactivated BMDM supernatants were prior treated for 30 minutes with different concentrations of anti-TNFα or anti-IFNAR and then added to the L929 cells as described above. For JAKi experiments, L929 cells were prior treated for 30 minutes with the specific inhibitor and then exposed to 50ul of the serially diluted UV inactivated BMDM supernatant, as described above. After 1-hour incubation –to test BMDM supernatant antiviral activity–cells were infected by adding 50ul of virus, at the desired MOI, in DMEM supplemented with 3% FCS, or left uninfected –to test the BMDM supernatant cytotoxic effect–by adding 50 µl of DMEM supplemented with 3% FCS without virus. 48 hours later, the vitality of the cells was evaluated by CellTiter-Glo luminescent cell viability assays (Promega, Madison, WI, USA), following the manufacturer’s instructions.

### Viral titration (TCID50)

Titration for MHV and SARS-CoV2 was performed using the TCID50 method, with some specifications. Briefly, supernatants from infected cells were 10-fold serially diluted and titrated on target cells plated at 80-90% confluence (for MHV, L929 cells; for SARS-CoV2, VeroE6 cells) in 96 wells in a total volume of 100 µl, (8 replicate wells for MHV, 10 replicate wells for SARS-CoV2) and incubated at 37 °C under 5% carbon dioxide. After a defined time period (2 days for MHV, 5 days for SARS-CoV2), each well was scored for the presence of virus-induced cytopathic effects. The limiting dilution end point (TCID50) was determined using the Reed–Muench method for MHV and the Kärber equation for SARS-CoV2.

### Generation of LSD1 KO cells by CRISPR/Cas9

The sgRNA oligo (see Reagents table) ^10^, targeting exon 3 of Lsd1, was cloned into pSpCas9(BB)-2A-GFP, a gift from Feng Zhang (Addgene plasmid # 48138; http://n2t.net/addgene:48138; RRID:Addgene_48138) ^52^. The plasmid was subsequently transfected, in parallel with empty vector controls, in L929 cells using Lipofectamine 2000 according to manufacturer’s instructions. Forty-eight hours later, GFP-positive cells were single-cell sorted in 96-well plates and after clonal expansion, sublines were screened by Western blot against LSD1.

### RNAi Knockdown

To knock down LSD1, one short hairpin RNA (shRNA) sequences was tested. shLSD1#1 (see reagents table for sequence) was cloned into the pLKO vector by AgeI–Eco RI double digestion. The plasmid was used to produce lentiviral particles in 293T cells.

### Size Exclusion Chromatography

5 ml of UV-inactivated cell culture supernatants were fractionated into 53 fractions on a Superdex 200 16/60 column (Cytiva Life Sciences) with PBS as eluting buffer. 1ml fractions were collected and analyzed by antiviral and cytotoxic assays, and ELISA.

### Live cell imaging

L929 (20k cells), BMDM (5k or 10k cells), BMDM:L929 1:1 (12.5k:12.5k cells) and BMDM:L929 2:1 (20k:10k cells) cocultures were seeded on a 96-wells plate at day 0, treated at day1, infected with MHV 0.1MOI at day 2 for 1h, washed and kept in fresh medium plus treatments and 0.4ug/ml Propidium Iodide (PI) for the total duration of the time-lapse experiment. GFP, PI and bright field images were acquired on a Nikon Eclipse Ti microscope (Nikon Instruments S.p.A., Firenze, Italy) equipped with a xyz motorized stage, a Spectra X light engine (Lumencor, Beaverton, OR, USA), a Zyla 4.2 sCMOS camera (Andor, Oxford Instruments plc, Tubney Woods, Abingdon, Oxon OX13 5QX, UK) a multi-dichroic mirror and single emission filters (Semrock, Rochester, New York, USA). Large images made by four partially overlapping (2%) fields of view (FOV) were acquired for each well with a 2×2 camera binning every hour from 4 hpi to 48 hpi using a 10x, 0.3 NA objective lens. Temperature, CO2 and humidity was controlled by a microscope cage incubator (Okolab, Napoli, Italy). Bone marrow derived macrophages or L929 cells were seeded on a 12 glass-bottom wells (MatTek Corporation, Ashland, MA 01721, USA), coated with poly-D-Lysine 0.1% (w/v) in water (3×105 cells/well). After 24 hours cells were treated with DDP (10 µM) or DMSO and then infected with MHV for 12 hours. Cells were incubated with 1 µM of Lysosensor Green DND-189 (Thermo Fisher Scientific, Monza, Italy) for 90 min. and Hoechst 33342 (Euroclone S.p.A., Pero, Italy) for the last 30 min. Cells were washed and fresh medium was added. Labelled live cells were imaged at 37 °C and 5% CO2 on a Leica Thunder Imager system (Leica Microsystems GmbH, Wetzlar, Germany), equipped with a xy motorized stage, 5 LED sources, a DFC9000 GTC sCMOS camera, a multi-dichroic mirror and 4 emission filters. Ninety-nine images were acquired for each condition using a 63x 1.4NA oil immersion objective lens.

### Immunofluorescence

Cells were seeded on Poly-D-lysine coated slides (10^5^ cells/slide). After treatments cells were fixed with methanol at −20°C for 6 min, blocked with 5% donkey serum for 60 min. Slides were stained with primary antibodies diluted in 1% BSA in PBS (NF-kB and IRF1 1:400, NSP9 1:1000) for 90 min. Secondary anti-rabbit (A488) and anti-mouse (Cy3) were used at 1:200 for 1 hour. Nuclei were stained with DAPI 1:1000 for 20 min. Mowiol was used as mounting solution. Cells labelled with DAPI and anti-NF-kB/NSP9 or anti-IRF1/NSP9 antibodies were imaged with a 60x 1.4 NA oil immersion objective lens on a CSU-W1 Yokogawa Spinning Disk confocal system with a 50 μm pinhole disk mounted on an Eclipse Ti2 stative and equipped with a motorized xyz stage, 6 solid state lasers, a multi-dichroic mirror, single emission filters and a Prime BSI sCMOS camera (TELEDYNE PHOTOMETRICS, Tucson, AZ 85706, USA). Hundred FOV per condition were automatically acquired thanks to the JOBS application of the NIS software (Nikon). Briefly, for each of the 100 positions defined in the JOB, an autofocus routine using the DAPI channel was used to define the acquisition focal plane and the 3 channels corresponding to the 3 stainings were acquired.

### Bioinformatic analysis of imaging data

#### Syncytia and death events quantification

Time-lapse raw images were first corrected for uneven illumination (shading correction) and background (BG) variation in time thanks to the BaSiC Fiji/ImageJ plugin ^53^. The corrections were done in batch using a custom made ImageJ macro. Briefly, for each 4-channel .nd2 time series, the channels were split, BaSiC-corrected and saved as tiff sequence in a new folder.

A custom-made ImageJ macro was used to identify syncytia formed by infected L929^GFP^ cells. Briefly, for each condition and in batch, the GFP time series was BG-corrected using a rolling ball radius of 50 pixels, filtered with a median filter (radius=2 pixels), then objects were identified using a threshold defined by the Otsu method, the objects with a size bigger than 10 um^2^ were identified by the Analyze Particles ImageJ function and the region of interest (ROI) added to the ROI manager. In each Field of View (FOV), the list of ROI was used to calculate the ROI area and the intensity of the PI channel.

The mean area of single nuclei was calculated from the control condition (DMSO, not infected) at the first time point and the mean nuclei area resulted about 150 um^2^. Syncytia were arbitrarily considered as the union of at least 5 nuclei, using an object size threshold of 750 um^2^. This threshold was visually confirmed to accurately capture the majority of syncytia.

To calculate the death events of L929 cells, the mean PI intensity in each ROI obtained from the segmentation of the GFP channel was considered, and an intensity threshold of 2000 grey levels was used to define a PI-positive object. For both populations, the number of objects were transformed in nuclei number (“death events”) multiplying every single object by a integer factor (>=1) calculated dividing the area of each identified object by the mean nucleus area (150 um^2^) and rounding to the minor integer. This correction was necessary to avoid underestimating death events associated with syncytia in-L929 infected cells. Live cells were calculated by dividing the total estimated nuclei by the number of death events, per each frame. In all figures, death events are expressed in absolute terms, whereas live cells are expressed relative to the number of live cells at the earliest recorded time frame (4 hpi).

### NFkB and IRF1 immunofluorescence quantification

Confocal images of BMDM were analyzed by a custom-made ImageJ macro written in Jython. Briefly, the DAPI channel, after a median filter and BG subtraction, was used to automatically segment the nuclei. To the nuclei binary image a Voronoi filter was applied to roughly identify the area relative to each cell. For each cell identified by a nucleus, a band around the nucleus with a thickness of 6 μm the cytoplasm was created and the intersection between the band and the cell area was considered as the “cytoplasm”. The total NSP9 intensity inside the cytoplasm was measured for each cell. The signal corresponding to 2 standard deviations above the mean NSP9 signal in the non-infected cells was used as threshold to identify the NSP9 positive and NSP9 negative cell populations in infected cells.

The IRF1/NF-κB nuclear signal was quantified inside the ROI obtained from the DAPI segmentation as mean intensity.

### Lysosensor assay

Bone marrow derived macrophages or L929 cells were seeded on a 12 glass-bottom wells (MatTek Corporation, Ashland, MA 01721, USA), coated with poly-D-Lysine 0.1% (w/v) in water (3×10^5^ cells/well). After 24 hours cells were treated with DDP (10 μM) or DMSO and then infected with MHV for 12 hours. Cells were incubated with 1 µM of Lysosensor Green DND-189 (Thermo Fisher Scientific, Monza, Italy) for 90 min. and Hoechst 33342 (Euroclone S.p.A., Pero, Italy) for the last 30 min. Cells were washed and fresh medium was added. Labelled live cells were imaged at 37 °C and 5% CO_2_ on a Leica Thunder Imager system (Leica Microsystems GmbH, Wetzlar, Germany), equipped with a xy motorized stage, 5 LED sources, a DFC9000 GTC sCMOS camera, a multi-dichroic mirror and 4 emission filters. Ninety-nine images were acquired for each condition using a 63x 1.4NA oil immersion objective lens. Widefield images of the DAPI and Lysosensor were quantified using a custom-made ImageJ macro. Similarly to what was done for the NSP9 quantification, the “cytoplasmic” Lysosensor total signal was calculated in a 6 μm thick band around the nucleus after BG subtraction.

### ChIP assay

Plates containing 15×10^6^ cells were washed 3 times with PBS and fixed at RT with 1% formaldehyde for 15 min. Cells were washed again 3 times with PBS, harvested with a cell lifter, collected into Falcon tubes and centrifuged at 424 rcf for 5 min at 4°C. Each pellet was resuspended in 3 ml of Lysis Buffer 1 (50 mM Hepes-KOH, pH 7.5, 140 mM NaCl, 1 mM EDTA, 10% glycerol, 0.5% NP-40 and 0.25% Triton X-100) and incubated on ice for 10 minutes. Nuclei were pelleted at 1600 rcf for 5 min. at 4°C, washed with 3 ml of Lysis Buffer 2 (10 mM Tris-HCl, pH 8.0 5M, 200 mM NaCl, 1 mM EDTA and 0.5 mM EGTA) and incubated at RT for 10 minutes. Nuclei were pelleted again at 424 rcf for 5 minutes at 4°C and the nuclear membrane was disrupted with 1,5 ml of Lysis Buffer 3 (10 mM Tris-HCl, pH 8.0, 100 mM NaCl, 1 mM EDTA, 0.5 mM EGTA, 0.1% Na-Deoxycholate, 0.5% N-lauroylsarcosine and protease inhibitors). Chromatin fragmentation was performed by sonication (Bioruptor® Plus sonication device, 45-60 cycles, 30 seconds on/off, high power, at 4°C). Chromatin extracts containing DNA fragments with an average of 300 bp were then subjected to immunoprecipitation.

The immunoprecipitation was performed using magnetic Dynabeads Protein G. Beads were blocked with 0,5% BSA in PBS and then mixed with the different antibodies (15 µg antibody:100 µl beads ratio) and incubated overnight on a rotating platform at 4°C. 1% of Triton X-100 was added to the sonicated lysates and lysates were centrifuged in microfuge (8000g, 10 min. at 4C) to pellet debris. Supernatants containing chromatin were subjected to immunoprecipitation and the 10% of the volume was used as input. Antibody-coated beads were added to each lysate and incubated overnight on a rotating platform at 4°C. Beads were washed 6 times (5 min each) with Wash Buffer (RIPA) and once with TE and 50 mM NaCl. The immunocomplexes were eluted in 100 µl of elution buffer (TE and 2% SDS) at 65°C for 15 minutes. To reverse cross-links immunocomplexes (and inputs) were treated with RNAase (0.5 mg/ml) at 37°C, 20 min followed by proteinase K and 1% SDS at 65°C overnight. DNA was purified using AMPure XP beads following manufacturer’s instructions. Isolated DNA was used to analyze by quantitative PCR the expression of NF-κB targets (TNFα, IL1β, CXCL1, CCL5, CXCL10 and a negative control), with the Fast SYBR™ Green Master Mix on a thermocycler Viia7 (Life Technologies, Inc.)

### RT-qPCR

Total RNA was purified using the RNeasy kit (Qiagen). RT-qPCR was performed using Luna® Universal One-Step RT-qPCR Kit (New England Biolabs), following the manufacturer’s instructions. Primers details are specified in reagents table.

### RNA sequencing

mRNA-seq libraries were prepared according to the TruSeq low sample protocol (Illumina, San Diego, CA, USA), starting with 1 µg of total RNA per sample. RNA-seq libraries were pair-end sequenced on an Illumina NovaSeq 6000 sequencing platform. RNA-seq data were mapped using STAR aligner ^54^ against the mouse genome (mm10). Counts were obtained by htseq-counts ^55^ and differential expression analysis was performed with DESeq2 package hosted in Galaxy online platform ^56^ using a false discovery rate (FDR) cut-off of 1 x 10^-4 9^. Hierarchical clustering was performed on z-score across samples for each gene, using Ward’s criterion with 1 - (correlation coefficient) as a distance measure.

### Transposable Element quantification from RNAseq

In order to quantify the expression of Transposable Elements (TEs) while avoiding the biases introduced by multimapping reads, we applied a clustering procedure that groups together TEs whose expression is supported by the same set of multimapping reads. Briefly, reads were mapped to the mouse reference genome (GRCm38 assembly) using Star and allowing an unlimited number of mapping locations. Then, BAM files for all samples were pooled and the coordinates of mapped reads were intersected (bedtools intersect ^57^) with those of TEs annotated in RepeatMasker (v405). This operation allowed us to build a binary matrix associating each read to the TEs that it maps to. Such matrix was then transformed with mcxarray into a square matrix of Tanimoto distances between each pair of TEs and subjected to clustering using the Markov Cluster Algorithm (MCL, filtering parameter >=0.5. Tanimoto distance. Inflation parameter 1.2 ^58^). This operation generated 514.866 clusters that contain a variable number of TEs with common mappability profiles. The number of reads in each cluster was then quantified for each sample (discarding reads mapping to multiple clusters) and the resulting TE expression matrix was imported into R for differential expression analysis with DESeq2 ^59^. Cluster counts for each sample were first normalised using size factors estimated from the number of reads uniquely mapping to Ensembl genes, then differential expression analysis was performed using the Wald Test and a design formula capturing the interaction between DDP treatment and MHV infection status (∼Infection*Treatment). The p-values thus obtained were then corrected for multiple hypothesis testing using the Benjamini-Hochberg procedure. To generate fold change boxplots for LINEs, SINEs and LTR clusters, we first discarded all clusters containing TEs belonging to multiple Repeat Masker classes. For the remaining TE clusters, we then plotted the log2 fold changes estimated by DESeq2 as boxplots. In order to test for differences in thesedistribution we used Welch’s t *test*.

### Viral genome analysis and assembly

To identify the MHV viral strain we used by the SPADES software ^60^. The longest contig obtained was compared with the most common strains of MHV (MHV1, MHV3, JHM, A59 - Supplementary Table 1) using QUAST ^61^.

### Western Blotting

For total protein lysates, around 5×10^6^ cells were lysed on the plate placed on ice, using 300 µl of RIPA buffer, containing the cOmplete™ Protease Inhibitor Cocktail for all experiments, except for experiment in Supplementary Figure 8A, in which 8M Urea buffer was used at RT. Cytoplasmic-nuclear protein fractions were performed as previously described (Czerkies et al., 2018). Cells were scraped in cold PBS, with a cell lifter, and centrifuged at 1350 rpm, 5 min at 4°C. Extracellular membranes were lysed, adding 1 ml of a hypotonic buffer (20 mM HEPES pH 8.0, 0.2% NP-40, 1 mM EDTA, 1 mM DTT and protease inhibitors) and incubating on ice for 10 min. The supernatant obtained after centrifugation at 1700 × *g*, 5 min at 4 °C contained the cytoplasmic protein fraction; the pelleted nuclei were washed with the same buffer and centrifuged again. The nuclear content was extracted by adding 150 µl of nuclear fraction buffer (20 mM HEPES pH 8, 420 mM NaCl, 20% glycerol, 1 mM EDTA, 1 mM DTT and protease inhibitors), incubating on ice for 30 min and isolating the supernatant after centrifugation at 10,000 × g, 10 min at 4°C. For both total and cytoplasmic-nuclear fractions, proteins were quantified using the Pierce™ BCA Protein Assay Kit (Thermo Fisher Scientific). Equal amount of proteins was mixed with Laemmli buffer and analyzed by SDS-PAGE, using nitrocellulose membranes. After blocking, membranes were incubated over night with the different primary antibodies at 4°C and secondary antibodies for 1 hour at RT. Digital images were obtained using Clarity Western ECL Substrate and the ChemiDoc^TM^ MP Imaging System (Bio-Rad).

### ELISA

To measure levels of cytokines secreted by BMDMs in response to MHV infection, ELISA was performed using 24 hrs post-infection supernatant from both infected and uninfected BMDM cultures. Murine TNF*α* levels were measured from 10ul of supernatant using the Mouse TNF*α* ELISA Set II kit (BD Biosciences) while murine IFN*α* was measured from 100ul of supernatant using the Verikine-High Sensitivity mouse IFN*α* all subtypes ELISA kit (PBL Assay Science). ELISAs were performed in duplicate according to manufacturer’s protocols. Similarly, TNF*α* and pan-IFN*α* were measured in fractions from size exclusion chromatography using 100ul of fraction volume.

## Acknowledgments

We would like to thank dr Federica Facciotti for critical reading of the manuscript, dr Tiziana Bonaldi and dr Alessandro Cuomo for support with fractionation experiments, dr Chiara Soriani for assistance with imaging anlaysis, dr Mario Varasi at IFOM for assistance in compound synthesis, dr Riccardo Villa from IZSLER – Brescia for providing the MHV strain.

## Author contributions

Conceptualization: LM, SM, PGP, FS, RR, PM, BAD

Methodology and validation: all authors

Investigation: LM, SM, PGP, FS, RR, PM, BAD, RP, EG, DT, MR, IP, CG, GB

Formal analysis: SR, EB, LM, TL

Writing-Original draft: LM

Writing-Review & editing: all authors

## Supplementary Materials

### Supplementary Note

#### Description of the experimental system

To investigate cell dynamics following MHV infection, we characterized viral titer, cell viability and syncytia formation (a typical but cell type-specific coronavirus-associated cytopathic effect) ^62^ in L929 fibroblasts and bone marrow-derived macrophages (BMDM) in response to the MHV strain A59 (see supplementary table 1 for viral strain confirmation). In both cell types, infection was productive, as demonstrated by a similar viral titer at 24 hours post-infection (hpi) (supplementary figure 1A). Viability decreased at 48 hpi in both cell types, following comparable MOI-dependent kinetics (supplementary figure 1B); syncytia were consistently seen in L929 cells, but not in BMDMs (supplementary figure 1C).

To analyse cellular interactions between L929 fibroblasts and BMDMs, we set up a live-cell imaging system that allowed monitoring of cell death and syncytia formation of L929 and BMDM in mono- and co-cocultures. To distinguish L929 cells from BMDMs and to allow precise quantification of syncytia formation, L929 cells were engineered to express H2b-GFP (L929^GFP^), while cell death was monitored by adding Propidium Iodide (PI). L929^GFP^ and L929^WT^ cells were equally permissive for infection (supplementary figure 1D).

In L929^GFP^ monocultures, we observed a rapid increase in syncytia numbers (supplementary figure 2A), starting from ∼12 hpi with a plateau at ∼30 hpi (supplementary figure 2B), followed by a wave of cell death starting from 33 hpi (supplementary figure 2C,D and supplementary video 1). In co-cultures of BMDM and L929^GFP^ cells, L929^GFP^ syncytia formed with kinetics similar to that in monoculture (supplementary figure 2A,B), while cell death was significantly anticipated by ∼15 hours (supplementary figure 2C,D and supplementary video 2), when no MHV-induced cytopathic effect is detectable. In contrast to L929 monoculture, in which death was equally distributed between syncytia and mono-nucleated cells, in cocultures death occurred predominantly outside syncytia (supplementary figure 1E and supplementary video 1-2). Importantly, viral titers were not increased by the presence of BMDM within 24 hpi (supplementary figure 1A), ruling out an increase in MHV virulence as a cause for early fibroblast death and suggesting the existence of a macrophage-dependent cytotoxic activity.

To characterize secreted activities, supernatants from BMDM infected at different MOI (from 0.001 to 0.1) were collected 24 hpi, UV-inactivated, and tested in progressive 3-fold dilutions on L929 cells, which were infected or not with MHV at MOI 0.1 (see figure 1E for a schematic view of the experimental protocol). Supernatants from non-infected BMDMs had no effect on L929 cells, infected or not (supplementary figure 2F, I and supplementary figure 1F). Supernatants from infected BMDMs, in turn, showed complex effects: at high MOI (0.01-0.1) and high concentrations (1:3-1:9 dilution), they strongly reduced syncytia formation (supplementary figure 2G,K and supplementary video 3) and viral titer (supplementary figure 2H), but also induced early cell death, resulting in decreased overall viability (supplementary figure 2F). Death was MHV-independent, as it was equal in infected (figure 1J, K) and noninfected (supplementary figure 2L, supplementary figure 1F and supplementary video 6) L929 cells, and started immediately after supernatant addition, a kinetics incompatible with prior viral cytopathic effect. At intermediate dilution (1:27), both syncytia formation and early MHV-independent cell death were significantly reduced, resulting in maximal gain in overall viability of infected cells (figure 1I and supplementary videos 4 and 7). At further dilutions (1:81, 1:243), both antiviral activity (syncytia formation) and MHV-independent death were lost, giving way to late MHV-dependent death (figure 1J and supplementary videos 5 and 8).

These experiments suggested the existence of two biochemically distinct species of secreted activities with antiviral and cytotoxic effect, respectively. To verify this, we subjected the supernatants from infected macrophages to size-exclusion chromatography through gel filtration and measured the effects of the supernatant fractions on cell viability of MHV-infected or uninfected cells. We operatively defined the loss of cell viability in uninfected cells as “extrinsic cytotoxic activity” (ECT) and the rescue of cell viability in infected cells as “extrinsic antiviral activity” (EAV). As shown in figure 2A-C, the two activities eluted in two clearly separated size ranges, with ECT peaking in fractions 21-25 (corresponding to a predicted protein size of 50-60 kDa) and EAV peaking in fraction 27-31 (corresponding to a predicted protein size of ∼20 kDa).

Previous studies identified TNF*α* and type I interferon as candidates for coronavirus-induced cytotoxic and antiviral activities respectively^25, 29^. ELISA showed elevated levels of TNF*α* and IFN*α* in the fractions showing maximal ECT and EAV, respectively (figure 2B,D), consistent with their expected size (TNF*α* is biologically active in 52 kDa trimers ^24^. Neutralization experiments confirmed the role of TNF*α* and IFN*α*, since: i) anti-TNF*α* antibodies dose-dependently rescued viability of both infected and uninfected cells (figure 2E); ii) anti-IFNAR antibodies (figure 2F) or JAK2 inhibitors (which block IFN signal transmission, supplementary figure 2A) had no effect on ECT (viability in uninfected cells remained low) and antiviral activity could not be adequately measured due to widespread cell death; iii) dual IFNAR + TNF*α* blockade resulted in loss of both activities: viability was fully restored in uninfected cells (figure 2G), but syncytia became evident in infected cells (figure 2H).

These results unequivocally demonstrate that coronavirus-infected macrophages secrete biochemically separated cytotoxic and antiviral activities, identified as TNF*α* and IFN*α* respectively. Additionally, supernatant transfer provides an easily tractable model to independently quantitate macrophage extrinsic activities, potentially useful for genetic or pharmaceutical screens.

---

**Fig. S1.**
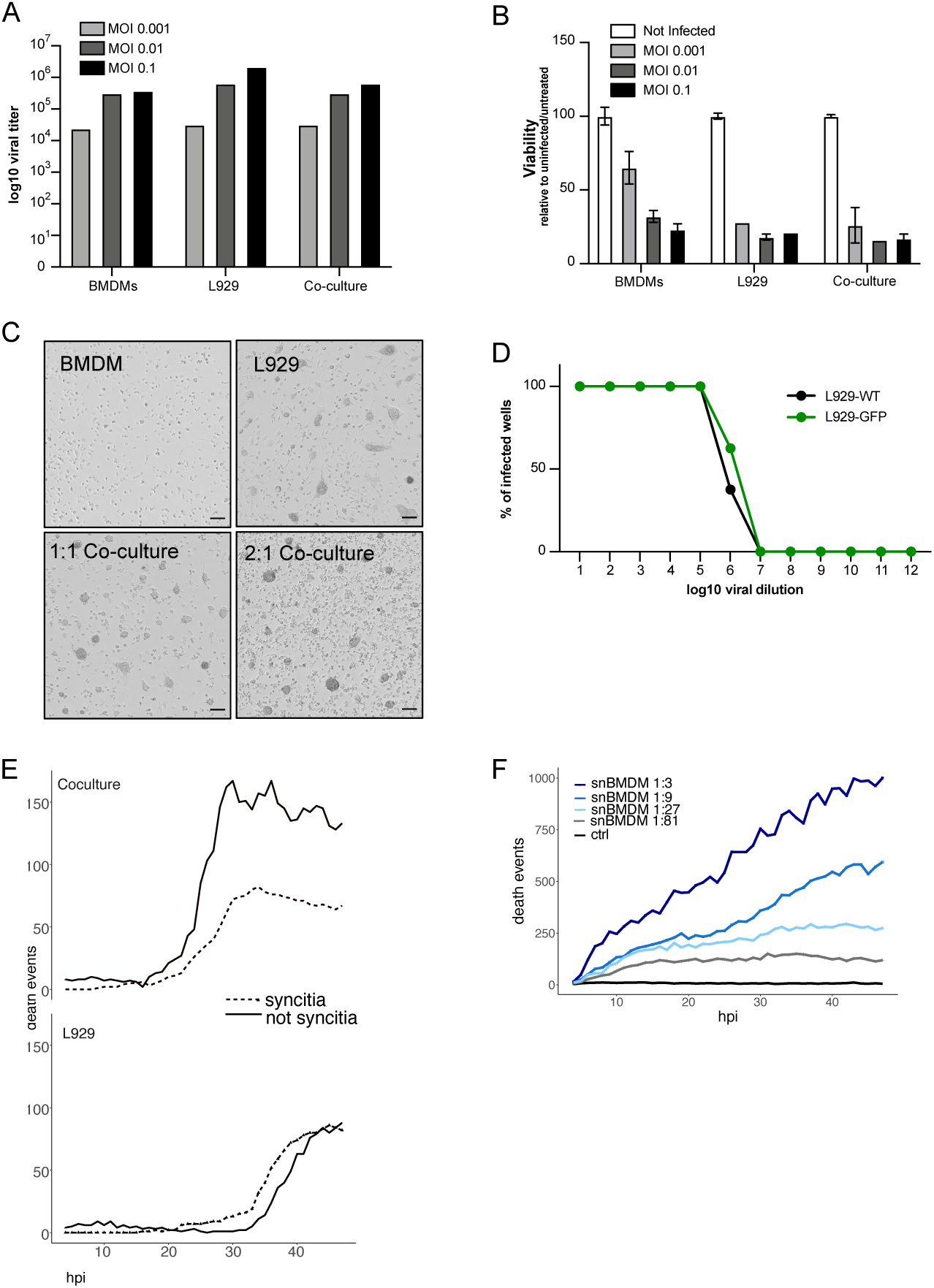
**A.** Viral titer 24 hpi in BMDM, L929 and 2:1 cocultures, by MOI, measured by TCID50 **B.** Cell viability 48 hpi measured by CTG in BMDM, L929 and 2:1 cocultures, by MOI. Mean and standard deviation of triplicate wells, expressed as percentage relative to uninfected cells **C.** representative bright field images of mono- and cocultures 48 hpi. Scale bars: 100 μm. **D.** TCID50 of L929^GFP^ vs L929^WT^ cells **E.** Death events overlapping with syncytia or non-syncytia in L929^GFP^ monocultures (upper panel) or 2:1 BMDM:L929 cocultures (bottom panel) **F.** Time course of death events in time lapse microscopy, of uninfected L929^GFP^ cells (MOI 0.1) exposed to the supernatant of MHV infected BMDM (MOI 0.1) at the indicated dilutions. Normal medium was used as control (CTRL).

**Fig. S2.**
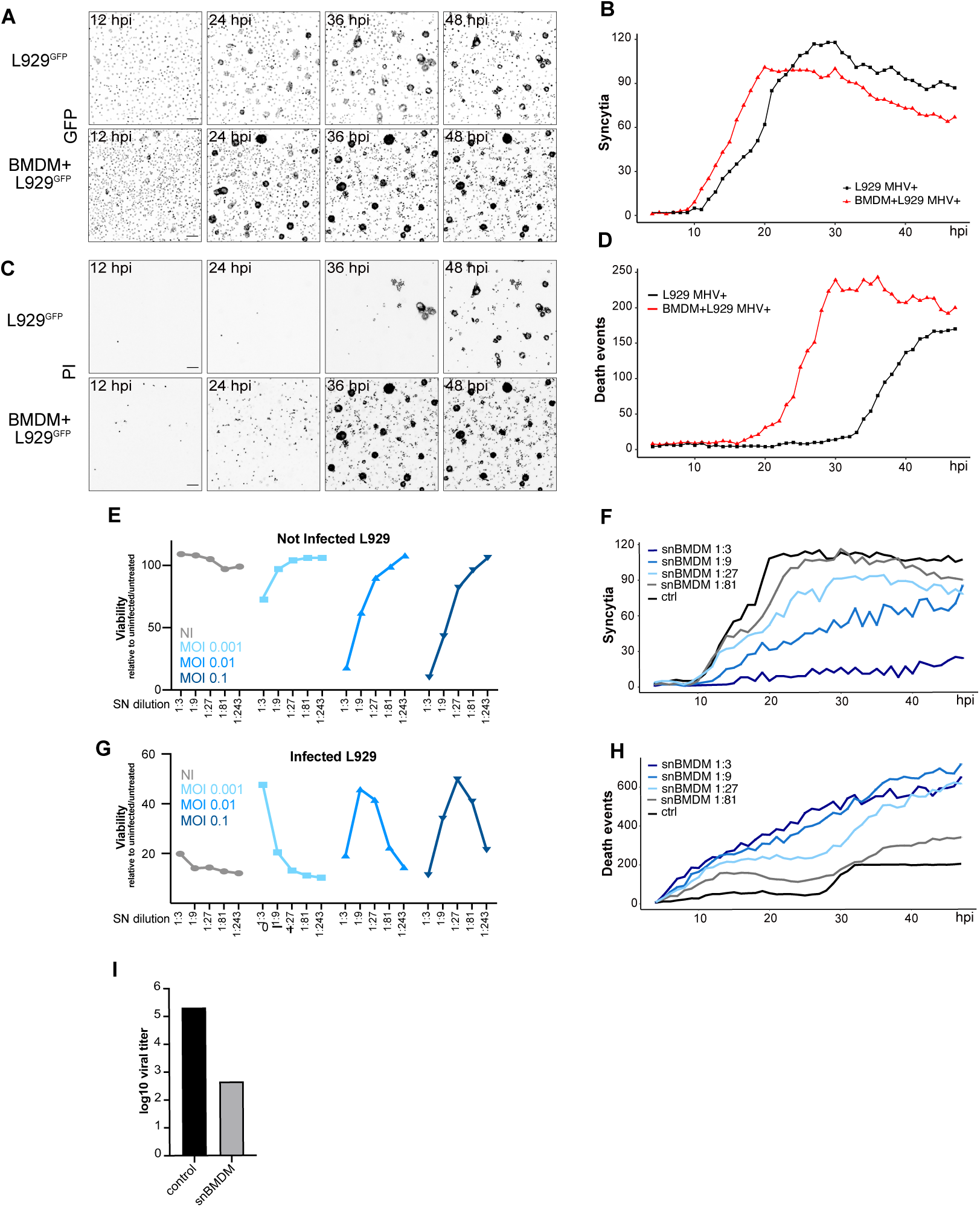
**A, C:** Representative snapshots from time lapse microscopy imaging of L929^GFP^ cells alone or cocultured with BMDM at a 1:2 ratio. Cells were infected at MOI 0,1. hpi: hours post infection. A, GFP signal, C: PI signal (cell death). Fluorescence signals were visualized with the inverted grey look up table (LUT). Scale bars: 100 μm. **B, D**. Time course of syncytia formation (B) and death events occurrence (D), monitored by time lapse microscopy, of L929^GFP^ cells alone or cocultured with BMDM at a ratio 1:2 respectively. Cells were infected at MOI 0,1. hpi: hours post infection. **F,I:** Cell viability of uninfected (F) and infected (I) L929 cells exposed to BMDM supernatants at three-fold dilutions (from 1:3 to 1:243). BMDM supernatants were collected from not infected (NI) or MHV infected BMDM at different MOI (MOI 0,1-0,001). Viability was measured by Cell Titer Glo in triplicate and expressed as Relative Luminescence Units (RLU) compared to uninfected and untreated cells. **G,J**: Time course of syncytia formation (G) and death events occurrence (J), monitored by time lapse microscopy, of MHV-infected L929^GFP^ cells (MOI 0.1) exposed to the supernatant of MHV infected BMDM (MOI 0.1) at the indicated dilutions. Normal medium was used as control (CTRL). **H:** viral titer from L929 cells treated (sn BMDM) or not (control) with supernatant from MHV-infected BMDM (MOI 0,1) at 1:3 dilution. Viral titers were measured by TCID50, 24 hours post infection.

**Fig. S3.**
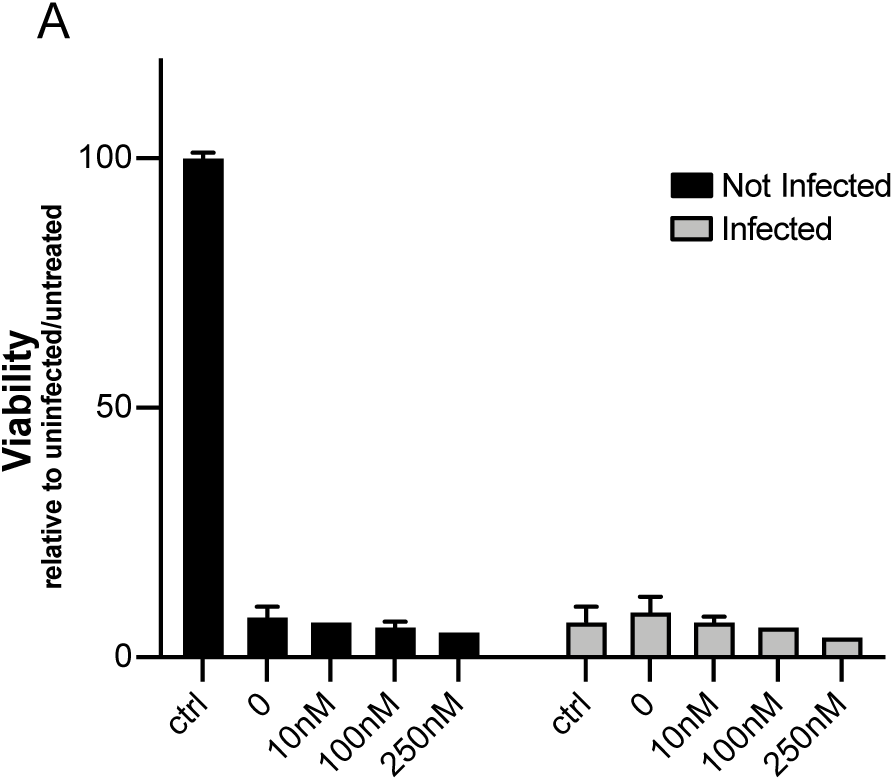
A. Effect of increasing doses of JAK inhibitor I, on 48h viability of non-infected or infected MOI 0.1 L929 cells. Mean ±SD of triplicate wells per experiment

**Fig S4.**
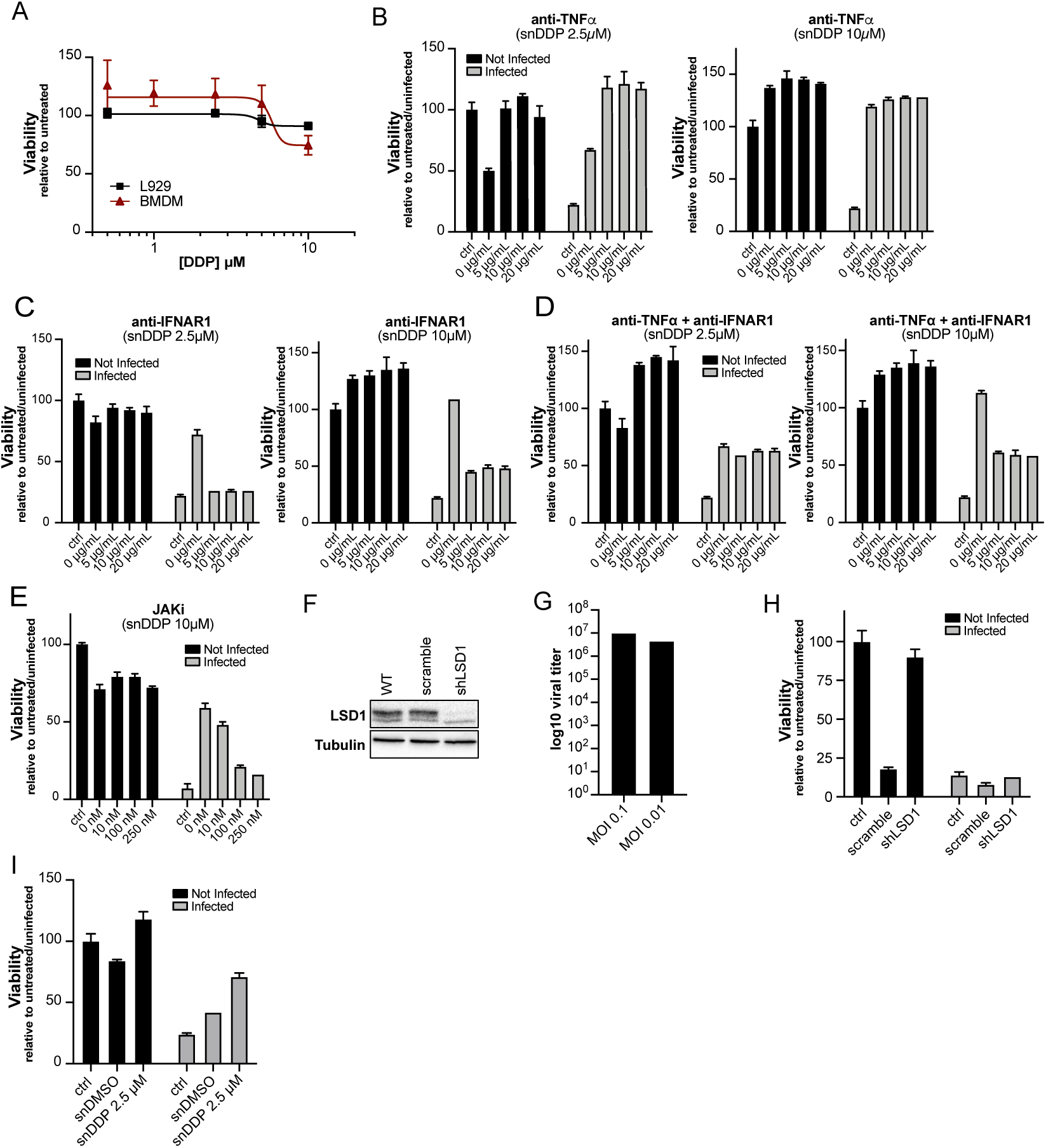
**A.** Effect of DDP38003 on viability of L929 and BMDM cells **B-E**. Effect of increasing doses of anti-TNF*α* (B), anti-IFN*α* (C), anti-TNF*α* + anti-IFN*α* (D) or JAK inhibitor I (E), on 48h viability of non-infected or infected MOI 0.1 L929 cells exposed to supernatant from MOI 0.1-infected BMDM treated with DDP 2.5 or 10 μM. Mean ±SD of triplicate wells per experiment **F.** Efficiency of shLSD1 on LSD1 protein levels in RAW264.7 cells **G.** Viral titer 24 hpi with MHV-A59 MOI 0.1 in Raw 264.7 cells, measured by TCID50 **H.** Viability of infected (MOI 0.1) or uninfected L929 cells exposed for 48 hours to empty medium (control) or snBMDM scramble and shLSD1 infected RAW264.7 cells **I.** Viability of infected (MOI 0.1) or uninfected LA4 cells exposed for 48 hours to empty medium (control) or supernatant from BMDM MOI 0.1 infected and treated with DMSO or DDP 2.5 μM

**Fig S5.**
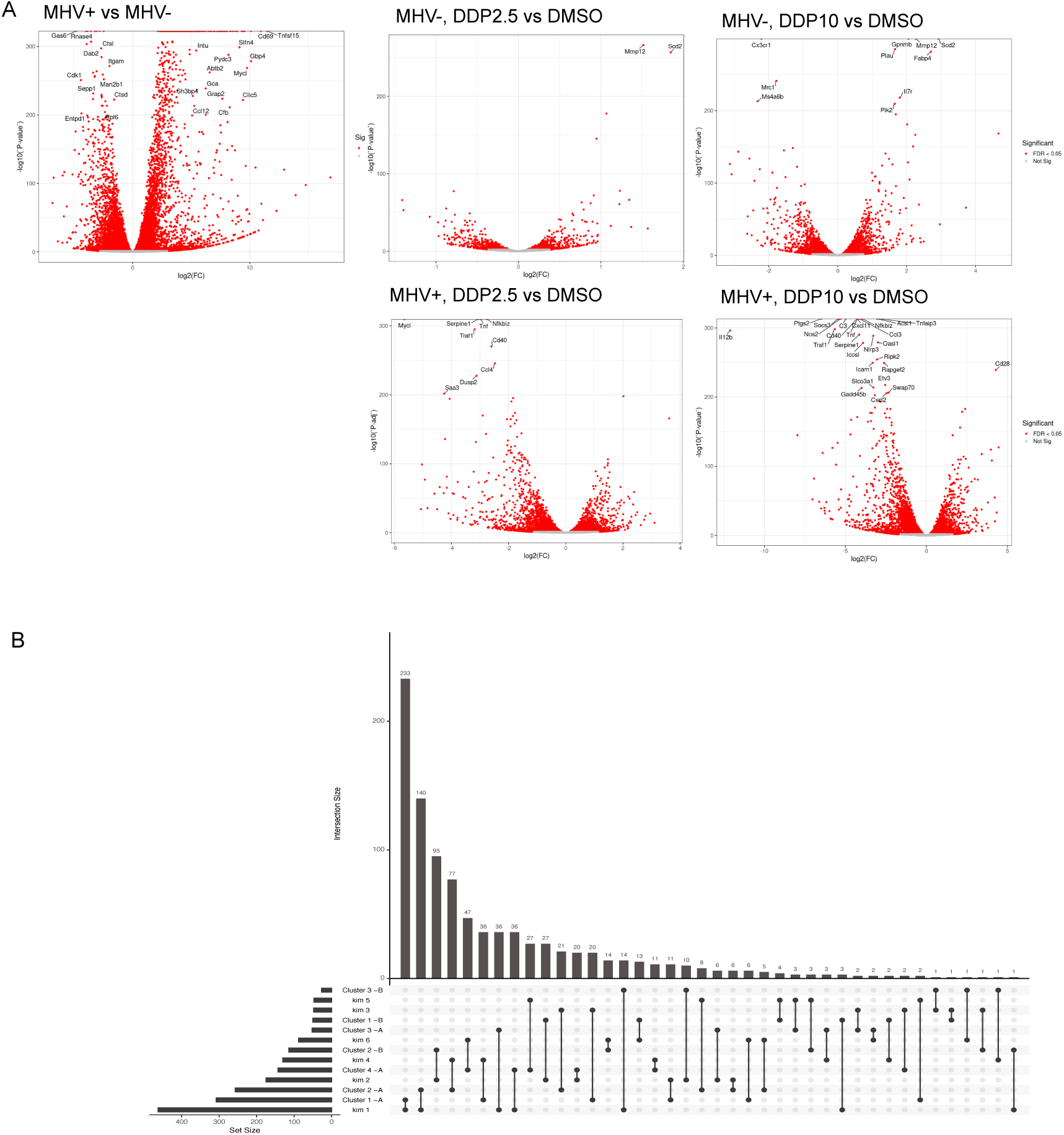
**A.** Volcano plots of indicated contrasts in BMDM infected or not with MHV-A59 MOI 0.1 at 24 hpi and treated with vehicle (DMSO) or DDP at 2.5 or 10 μM **B.** Upset plot showing overlap between genes in BMDM clusters and clusters in Kim et al ^9^

**Fig S6.**
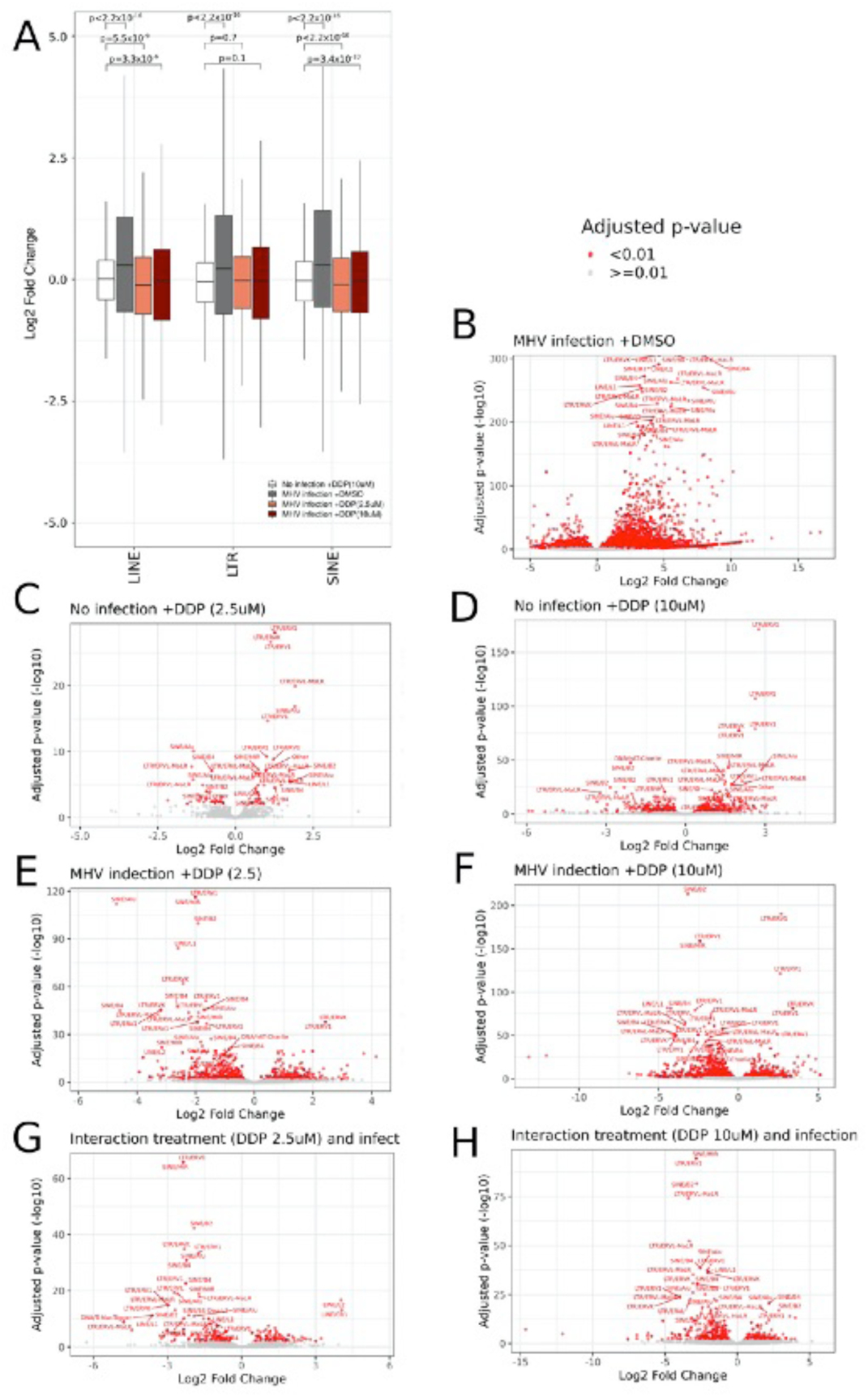
**A.** Boxplot showing the distribution of log2 Fold Changes for LINEs, LTRs and SINEs upon DDP treatment in non-infected BMDM cells (vs DMSO treated cells, white), untreated MHV infected cells (vs DMSO treated, uninfected cells, grey), MHV infected cells treated with DDP 2.5uM (vs MHV infected cells treated with DMSO, salmon) and MHV infected cells treated with DDP 10uM (vs MHV infected cells treated with DMSO, red). The p-values reported were obtained with Welch’s t-test. **B-F**. Volcano plot showing differential expression results for TE clusters (see material and methods) for the same comparisons as in B. **G,H**. Volcano plot showing interaction coefficients for DDP treatment and MHV infection. The interaction coefficients (x-axis) correspond to the difference between the log2 Fold Change of DDP-treated, MHV infected cells (vs DMSO-treated, MHV infected cells) and the log2 Fold Change of DDP-treated, uninfected (vs DMSO-treated, uninfected cells).

**Figure S7.**
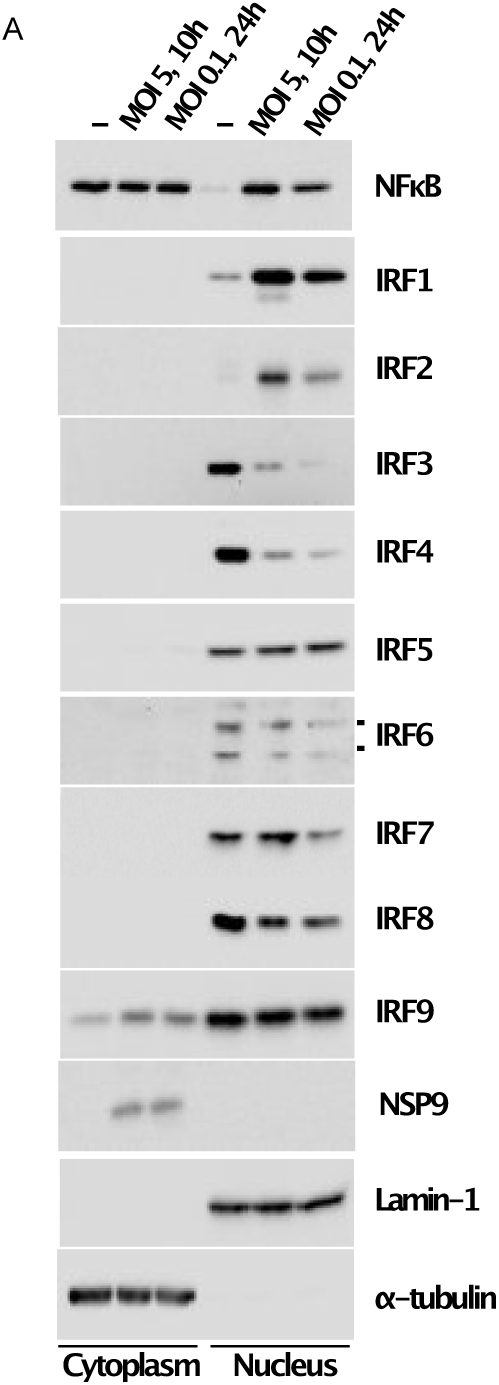
**A.** Expression of IRF factors in nuclear extracts of infected BMDM at 8h (MOI 5) and 24h (MOI 1), by Western Blotting

**Figure S8.**
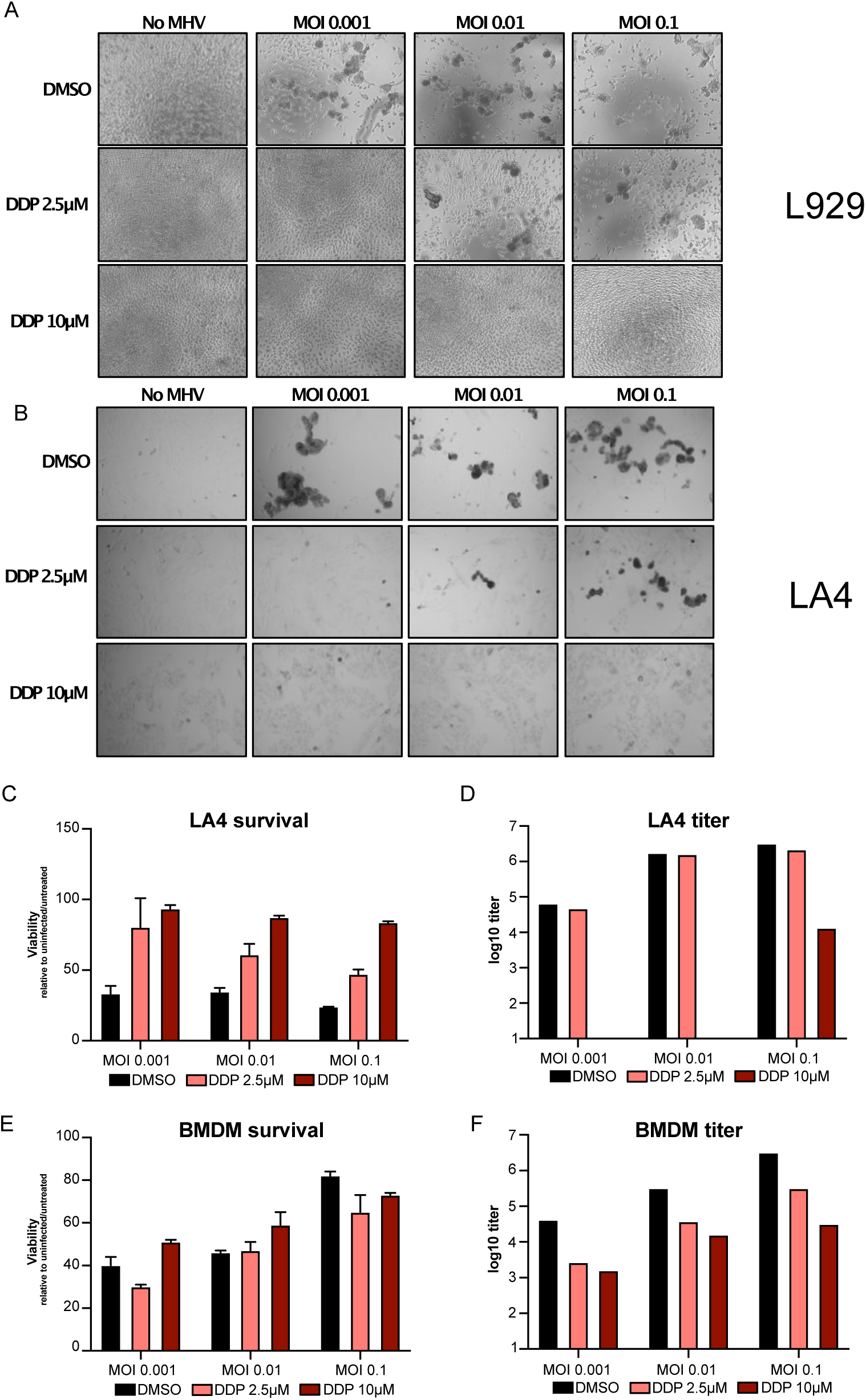
**A.** Bright field image of L929 cells infected with MHV-A59 and treated with DDP at indicated doses and MOI **B.** Bright field image of LA4 cells infected with MHV-A59 and treated with DDP at indicated doses and MOI **C.** Survival (measured by Cell Titer Glow) of LA4 cells infected with MHV-A59 and treated with DDP at indicated doses. Mean ±SD of triplicate wells per experiment **D.** Viral titer (TCID50) of LA4 cells infected with MHV-A59 and treated with DDP at indicated doses **E.** Survival (measured by Cell Titer Glow) of BMDM infected with MHV-A59 and treated with DDP at indicated doses. Mean ±SD of triplicate wells per experiment **F.** Viral titer (TCID50) of BMDM infected with MHV-A59 and treated with DDP at indicated doses and MOI

**Figure S9.**
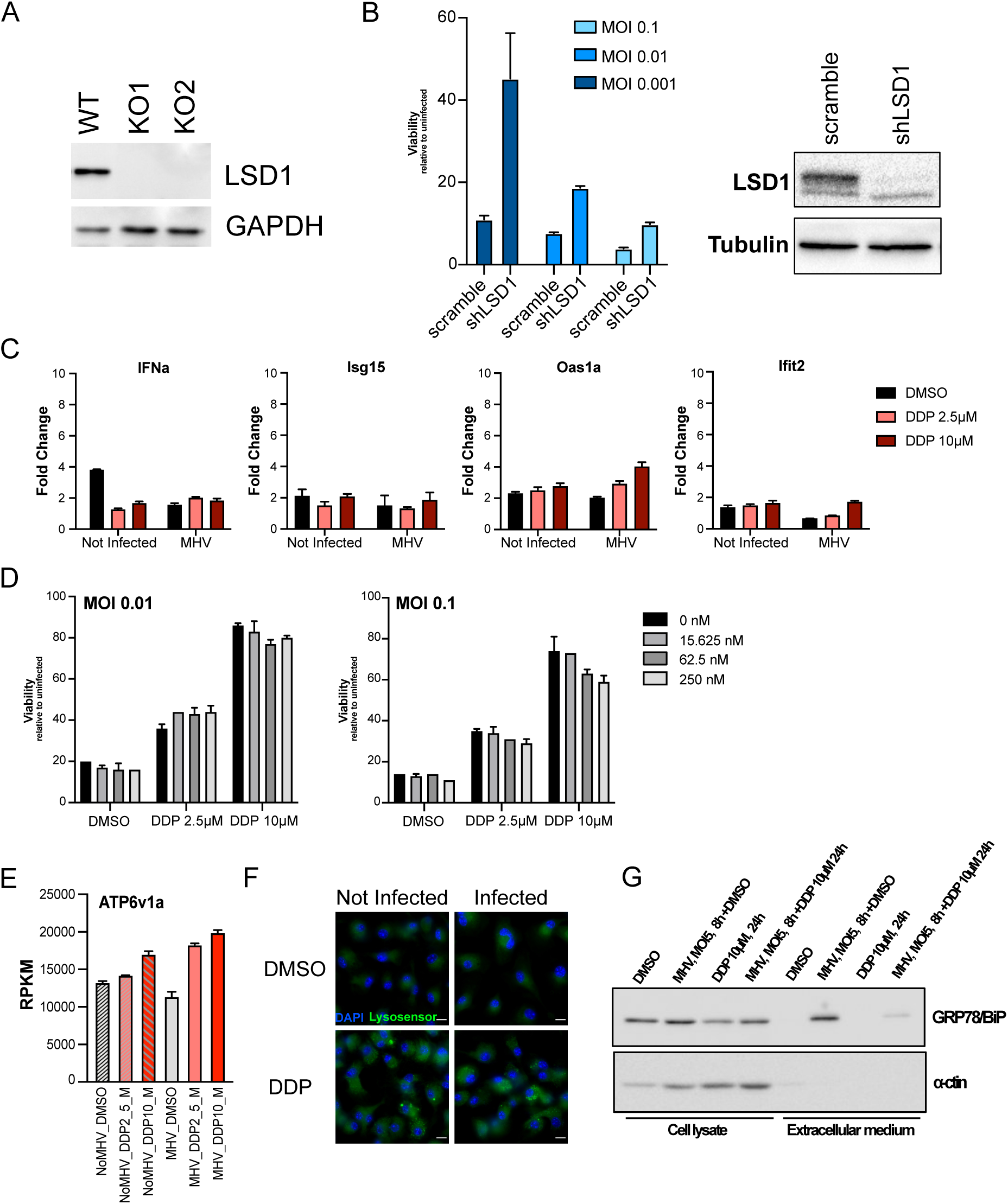
**A.** Western Blot showing ablation of LSD1 protein in L929 cells by CRISPR-Cas9 **B.** Effect of LSD1 knockdown on L929 cell survival measured by CTG after infection with different MOIs of MHV-A59. Mean ±SD of triplicate wells per experiment. On the right, WB demonstrating efficient though incomplete LSD1 knockdown **C.** Expression of IFN*α* and selected ISGs by qRT-PCR in response to MHV-A59 MOI 0.1 24 hpi and DMSO or DDP at 2.5 or 10 μM. Values are expressed as fold change over Time 0. Mean±SD of triplicate wells **D.** No effect of JAK inhibitor I on DDP rescue of cell viability of infected L929 cells at different MOI as indicated. Mean ±SD of triplicate wells per experiment **E.** Expression of the lysosomal ATPase in BMDM (RPKM from RNAseq in figure 4) **F.** Lysosome pH measurement using lysosensor on BMDM uninfected or infected with MHV-A59 MOI 5 and treated with DMSO or DDP 10 μM, at 12 hp **G.** BiP extrusion upon infection on BMDM uninfected or infected with MHV-A59 MOI 5 and treated with DMSO or DDP 10 μM, at 12 hpi, measured by WB on cell pellets and 1000 μl supernatant).

**Fig S10.**
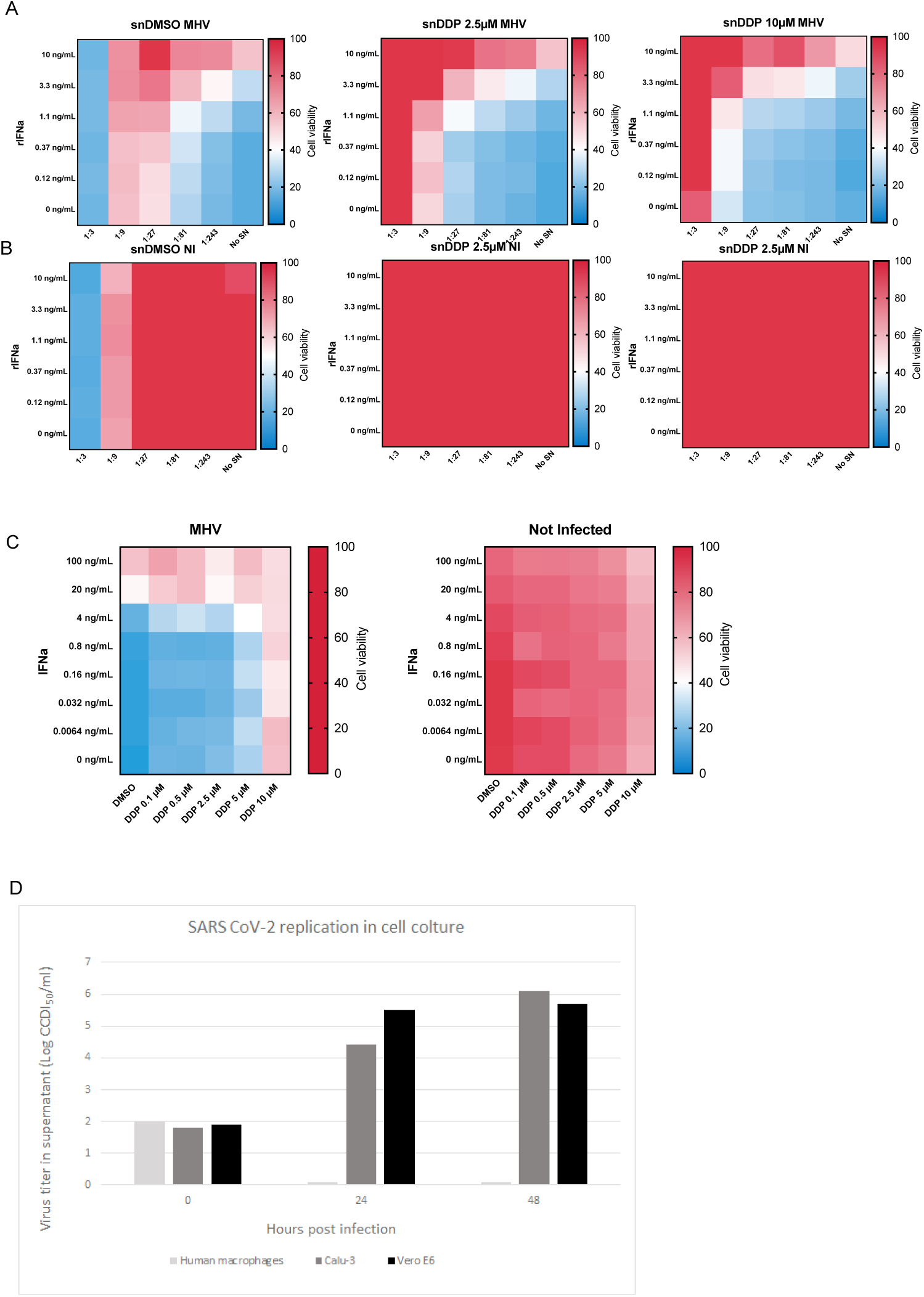
**A.** Cell viability by CTG (relative to uninfected) of infected L929 cells co-treated with with IFN*α* and supernatant of infected BMDMs treated with DMSO or DDP 2.5 or 10 μM **B.** Cell viability by CTG (relative to uninfected) of uninfected L929 cells co-treated with with IFN*α* and supernatant of infected BMDMs treated with DMSO or DDP 2.5 or 10 μM **C.** Cell viability by CTG (relative to uninfected) of infected L929 cells co-treated with with IFN*α* and DMSO or DDP 2.5 or 10 μM **D.** Viral titer in human cells (CD14-derived macrophages, Calu3 and VeroE6) in response to in vitro inoculation of SARS-CoV2 MOI 0.1, showing lack of productive infection in macrophages

**Movie S1.**Time lapse microscopy of L929^GFP^ cells infected with MHV-A59 at MOI 0.1. One frame per hour from 4 to 47 hpi. Brightfield, GFP (in green) and PI (in red) channels are overlaid. Scale bar: 100 μm.

**Movie S2.** Time lapse microscopy of 2:1 BMDM:L929^GFP^ cells infected with MHV-A59 at MOI 0.1. One frame per hour from 4 to 47 hpi. Brightfield, GFP (in green) and PI (in red) channels are overlaid. Scale bar: 100 μm.

**Movie S3.** Time lapse microscopy of L929^GFP^ cells infected with MHV-A59 at MOI 0.1 and treated with BMDM supernatant at 1:3 dilution. One frame per hour from 4 to 47 hpi. Brightfield, GFP (in green) and PI (in red) channels are overlaid. Scale bar: 100 μm.

**Movie S4.**Time lapse microscopy of L929^GFP^ cells infected with MHV-A59 at MOI 0.1 and treated with BMDM supernatant at 1:27 dilution. One frame per hour from 4 to 47 hpi. Brightfield, GFP (in green) and PI (in red) channels are overlaid. Scale bar: 100 μm.

**Movie S5.** Time lapse microscopy of L929^GFP^ cells infected with MHV-A59 at MOI 0.1 and treated with BMDM supernatant at 1:243 dilution. One frame per hour from 4 to 47 hpi. Brightfield, GFP (in green) and PI (in red) channels are overlaid. Scale bar: 100 μm.

**Movie S6.** Time lapse microscopy of uninfected L929^GFP^ cells and treated with BMDM supernatant at 1:3 dilution. One frame per hour from 4 to 47 hpi. Brightfield, GFP (in green) and PI (in red) channels are overlaid. Scale bar: 100 μm.

**Movie S7.** Time lapse microscopy of uninfected L929^GFP^ cells and treated with BMDM supernatant at 1:27 dilution. One frame per hour from 4 to 47 hpi. Brightfield, GFP (in green) and PI (in red) channels are overlaid. Scale bar: 100 μm.

**Movie S8.**Time lapse microscopy of uninfected L929^GFP^ cells and treated with BMDM supernatant at 1:243 dilution. One frame per hour from 4 to 47 hpi. Brightfield, GFP (in green) and PI (in red) channels are overlaid. Scale bar: 100 μm.

